# Cooperation between cortical and cytoplasmic forces shapes planar 4-cell stage embryos

**DOI:** 10.1101/2025.11.24.690087

**Authors:** Silvia Caballero-Mancebo, Daniel Gonzalez Suarez, Janet Chenevert, Sameh Ben-Aicha, Lydia Besnardeau, Alex McDougall, Rémi Dumollard

**Affiliations:** Laboratoire de Biologie du Développement de Villefranche-sur-Mer, Institut de la Mer de Villefranche-sur-Mer, Sorbonne Université, CNRS, 06230 Villefranche-sur-Mer, France; Institut de Biologie de Valrose, Université Côte-d’Azur, CNRS, 03108 Nice, France

## Abstract

Early embryonic cleavages often follow conserved geometric rules, resulting in species-specific cleavage patterns. How these rules are mechanistically implemented, however, varies widely across species and remains poorly understood. Here, using quantitative 3D live imaging, mechanical and biochemical manipulations, we dissect the mechanisms governing centrosomal complex (CC) migration and spindle orientation in ascidian 2-cell stage embryos, which generate the characteristic planar, square 4-cell stage. We show that following the first mitotic division, the CCs in each blastomere form at variable orientations relative to the mother spindle and progressively achieve parallel and coplanar alignment during interphase of the 2-cell stage. Final CC orientation is established before nuclear envelope breakdown, contrasting with the dynamic spindle reorientation during mitosis reported in other organisms. Our analyses reveal that CC rotation is driven by forces acting on long astral microtubules and guided by an anisotropic, endoplasmic reticulum (ER)-rich domain that surrounds the CCs. This ER domain is associated with a dense astral microtubule network, enabling efficient length-dependent microtubule pulling that can orient the CCs even in the absence of cell shape cues. In parallel, dynein-mediated cortical pulling refines CCs tilt and maintains coplanarity. Together, these findings uncover a cell-cycle-regulated cooperation between ER-mediated cytoplasmic forces and anisotropic cortical forces that ensures robust planar cleavage in ascidian early embryos.

## INTRODUCTION

Species-specific patterns of early cleavage divisions have fascinated developmental biologists for more than a century. Despite the diversity of embryo sizes and shapes, early cleavages typically follow two conserved geometric principles: Hertwig’s rule, which states that the mitotic spindle aligns along the longest axis of the cell (Hertwig, 1884), and Sachs’ rule, where division occurs at right angles to the previous one (Sachs, 1877). These rules collectively explain the widespread emergence of the planar and square arrangement of blastomeres at the 4-cell stage, a configuration observed across a broad spectrum of animal taxa (Caballero-Mancebo et al., 2026; McDougall et al., 2019; Pierre et al., 2016). Yet, despite the apparent universality of these geometric rules, the mechanisms by which they are implemented are species-dependent and remain the subject of active investigation and debate.

The physical environment of the embryo can impose significant constraints on blastomere arrangement and cleavage orientation. Nematode embryos, encased in a rigid eggshell, display a striking diversity of 4-cell stage morphologies, ranging from square to linear configurations, depending on the mechanical properties and the aspect ratio of the eggshell (Giammona and Campàs, 2021; Schulze and Schierenberg, 2011; Seirin-Lee et al., 2022). In contrast, unconfined embryos and embryos with a non-restrictive confinement, such as ascidian and echinoderm embryos, more consistently exhibit the archetypal flat, square 4-cell configuration (Minc and Piel, 2012; Pierre et al., 2016). However, this configuration is far from the mechanically stable configuration of 4 blastomeres—a tetrahedron—but arises due to the short division times of early embryos (Giammona and Campàs, 2021), suggesting that oriented cell divisions underpin the flat and square shape of 4-cell stage embryos.

To generate a symmetric and flat 4-cell stage embryo, the first two cell divisions must be equal and at 90° to each other, and the mitotic spindles of each cell of the 2-cell stage embryo must be coplanar and parallel. In large cells like the blastomeres of *Xenopus* embryos, geometric constraints coupled to yolk stratification are sufficient to generate flat 4-cell stage embryos (Pierre et al., 2016a; Wühr et al., 2010). This model postulates simply that cytoplasmic pulling forces acting on astral microtubules must be proportional to their lengths (Kimura and Kimura, 2011a; Pierre et al., 2016). This is thought to be achieved by dynein-dependent pulling of cytoplasmic cargoes moving towards the minus end of microtubules. However, the nature of these cargoes remains to be established (Xie et al., 2022).

In smaller cells, dynein-dependent cortical pulling forces become more prominent. In cultured cells and in cells of the *Drosophila* notum epithelium, cortical pulling forces have been shown to be essential in aligning the spindle with the long axis of the interphasic cell (Anjur-Dietrich et al., 2024; Bosveld et al., 2016). Moreover, in the *Caenorhabditis elegans* early embryo, where cytoplasmic pulling and cortical pulling or pushing have all been hypothesized to play a role in spindle orientation (Bondaz et al., 2019; Grill et al., 2001; Grill and Hyman, 2005; Kimura and Kimura, 2011b; Middelkoop et al., 2024; Wu et al., 2024), cortical pulling has been shown to be essential in the alignment of the spindle to the long axis of the zygote (Wu et al., 2024) as well as to the short axis of the AB cell at the 2-cell stage, thus having an opposing action to the length-dependent cytoplasmic pulling forces exerted on astral microtubules (Middelkoop et al., 2024). Overall, these observations argue for a size-dependent mechanism to orient spindles, where large cells rely on cytoplasmic pulling forces, whereas in small cells, cortical pulling forces dominate, and point to the intriguing possibility of intermediate-sized cells integrating both mechanisms to implement stereotyped oriented cell divisions.

In this study, we analyze the centrosome and spindle movements underlying the flat and square shape of the ascidian *Phallusia mammillata* 4-cell stage embryos, which are of intermediate size between the large *Xenopus* blastomeres and the small ones of nematode embryos. We find that centrosomal complexes (CCs) align progressively with the long axis of the blastomeres during interphase when cortical pulling is strong. Laser ablation experiments reveal that the force for the orientation of CCs relies on cytoplasmic pulling of microtubules on the stratified endoplasmic reticulum (ER). Finally, we find that interfering with the balance between cortical and cytoplasmic pulling forces transforms planar 4-cell stage embryos into non-planar or non-square embryos, highlighting that a finely tuned interplay between cytoplasmic and cortical pulling controls the precise positioning of spindles and the cleavage pattern of early embryos of intermediate size.

## RESULTS

### Progressive parallel and coplanar alignment of the two centrosomal complexes at the 2-cell stage

In order to follow the movements of each centrosome of the centrosomal complexes (CCs) during the 2-cell stage, we performed 3D live imaging of embryos injected with EB3-3xVenus to label the centrosomes from mitosis of 1-cell stage to the beginning of cytokinesis of the 2-cell stage (Fig. 1A, movie 1). In the vegetal view, two distinct centrosomes can be observed at each pole of the spindle at anaphase of the 1-cell stage, coinciding with the beginning of cytokinesis of the first mitosis (Fig. 1A-inset, t = 2 min). We found that the CCs in the two forming daughter cells are not initially parallel to each other (in 60% of all analyzed embryos) and progressively align with each other during the interphase of the 2-cell stage. This alignment results in two parallel CCs at nuclear envelope breakdown (NEB) of the 2-cell stage (Fig. 1A, 16-20 min), which in turn gives rise to two parallel spindles at metaphase of the 2-cell stage (Fig. 1A, 20-26 min). To generate the characteristic flat and square geometry of the ascidian 4-cell stage embryo, the two spindles at the 2-cell stage must not only be parallel but also coplanar; a side view revealed that, indeed, the two CCs remain largely coplanar throughout CCs migration at the 2-cell stage (Fig. 1A, bottom row).

**Figure 1:**
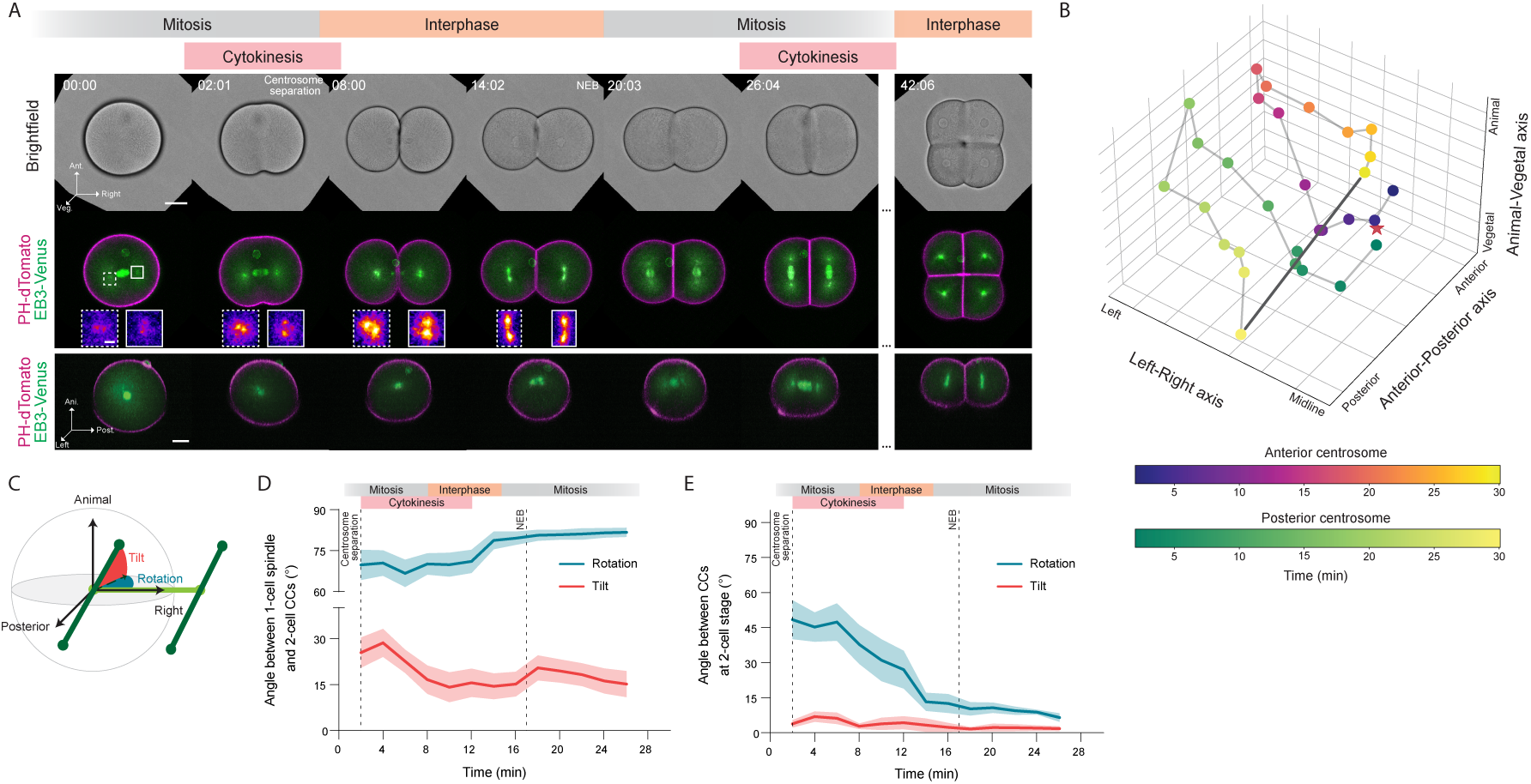
CCs become parallel and coplanar during interphase of the 2-cell stage. **(A)** Brightfield, confocal fluorescent images of a central z-slice of the plasma membrane (magenta), and maximum intensity projections of centrosomes (green) in an ascidian embryo from 1-cell stage metaphase (time = 00:00) to 4-cell stage interphase (time = 42:06). Top and middle rows show the vegetal view, and bottom row shows the side view. Insets are magnifications of the left (dashed square) and right (solid square) CC. Ant. is anterior, Veg. is vegetal, Ani. is animal, Post. is posterior, NEB is nuclear envelope breakdown. Time is in minutes:seconds. Scale bar, 30 µm. Inset scale bar, 5 µm. **(B)** 3D plot of the relative positions over time of the anterior and posterior centrosomes of the CC in the left cell of a 2-cell stage ascidian embryo. Time is coded in the respective color bar. The star represents the position of the left pole of the mitotic spindle at 1-cell stage, and the grey line represents the final orientation of the 2-cell stage spindle. **(C)** Graphical representation of the tilt and rotation angles in spherical coordinates between the CCs or the spindles at the 2-cell stage (dark green) and the spindle at the 1-cell stage (light green). **(D)** Plot of the rotation (blue) and tilt (red) angles between the 2-cell stage CCs or spindles and the 1-cell stage spindle as a function of time. The dashed line at 17 min indicates NEB. *N* = 9 embryos, *n* = 18 cells. Error bars, standard error of the mean. **(E)** Plot of the rotation (blue) and tilt (red) angles between the left and right CCs or spindles at 2-cell stage as a function of time. The dashed line at 17 min indicates NEB. *N* = 9 embryos. Error bars, standard error of the mean.

To further document the migration pattern of CCs and how they orient over time, we tracked the position of the anterior and posterior centrosomes within each CC in 3D (Fig. 1B) and measured the rotation and tilt angles of the segment formed by the centrosomes of each CC with respect to the mitotic spindle at 1-cell stage (Fig. 1C-D, S1A). We found that initially the CCs form with at a rotation angle of 68.2° ± 24.6 with respect to the mother spindle, and that during interphase of the 2-cell stage, the CCs rotate to achieve a final angle of 82.5° ± 5.3 with respect to the mitotic spindle at the 1-cell stage by the time the daughter cells are in metaphase of the 2-cell stage. Surprisingly, this final position of the CCs does not change much from the time of NEB, contrary to what has been observed in other embryos and somatic cells, where the final position of the spindle is fine-tuned during metaphase and anaphase (Anjur-Dietrich et al., 2024; Bondaz et al., 2019; Gönczy et al., 2001; Grill and Hyman, 2005), suggesting that spindle positioning in ascidian embryos is largely predetermined before NEB rather than dynamically adjusted during mitosis.

Consistent with this idea, the tilt angle between the CCs and the 1-cell stage spindle also decreased from 25.5° ± 22.4 to 15.1° ± 11.1 (Fig. 1D) and did not converge to zero. This observation is in agreement with previous reports describing a posterior-vegetal tilt of the spindle in ascidians at the 8-cell stage (Negishi et al., 2007). Comparing the orientation of the sister CCs and spindles at the 2-cell stage further confirmed our observations that they progressively become parallel, as indicated by a reduction in the rotation angle between them from 48.4° ± 25 to 6.6° ± 5.5 (Fig. 1E, S1B - Rotation), and that they are mostly coplanar throughout CC migration, as reflected by the small change in the tilt angle between them from 3.9° ± 4.8 to 1.8° ± 3.1 (Fig. 1E, S1B - Tilt). Altogether, this analysis revealed that the proper orientation of the CCs required to generate a flat, square 4-cell stage ascidian embryo is progressively established during the end of mitosis of the 1-cell stage and interphase of the 2-cell stage, and that by the time of mitotic entry at 2-cell stage, the final position of the future spindle is already defined.

To better understand the mechanisms that orient the CCs, we examined their migration at a higher spatiotemporal resolution (Fig. 2A, movie 2). We found that the CCs form in the center of the daughter cell at some distance from the nucleus (∼17 µm) before the first cleavage is complete (Fig. 2A, 30 s, arrowhead), and that the centrosomes within each CC remain close to each other and move together until the end of interphase of the 2-cell stage, when the centrosomes in each CC start to separate from each other and the nucleus positions between them (Fig. 2A-B, 12-14 min) right before NEB. Interestingly, the CCs have already reached their final orientation before their centrosomes start separating (Fig. 1D-E, 2B), indicating that the final orientation of the CCs is achieved without centrosome separation. Since the centrosomes within a CC remain close together during their migration, we asked whether a physical link between them is necessary for their proper migration. To that end, we performed periodic laser ablations every 90 s between the two centrosomes in one of the two sister CCs from the time when both centrosomes were identifiable (late anaphase of the 1-cell stage) until the nucleus reached the CC center (Fig. 2C, movie 3). We found that neither the rotation nor the tilt angles were affected when centrosomes were ectopically separated by laser ablations (Fig. 2C’, S2A-B), and that the ablated 2-cell stage embryos still divided into flat and square 4-cell stage embryos.

**Figure 2:**
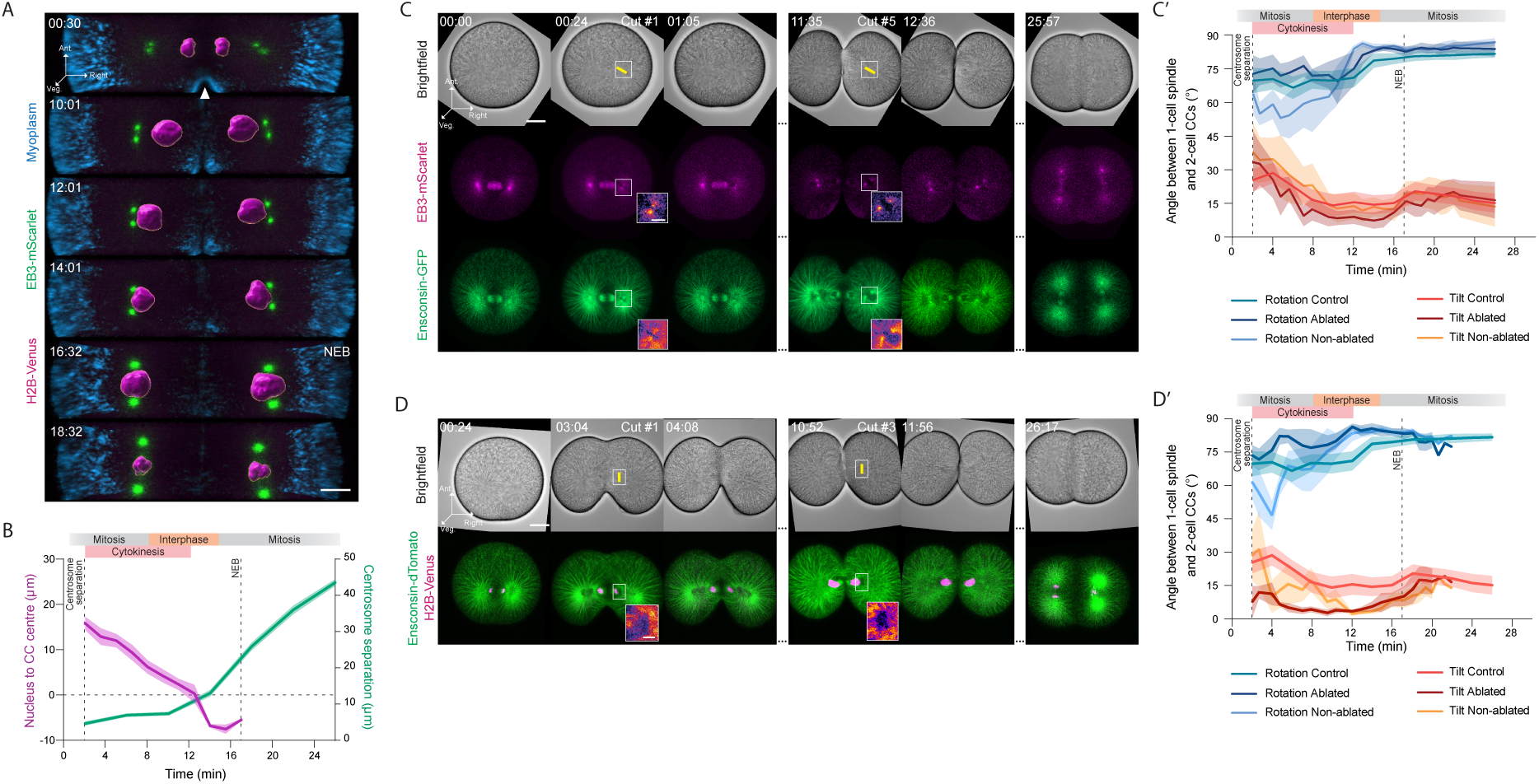
Migration of the CC and nucleus during interphase of the 2-cell stage. **(A)** Confocal maximum intensity projections of the centrosomes (green), mitochondria (cyan), and segmented nuclei (magenta) in an ascidian embryo during interphase of the 2-cell stage. Ant. is anterior, Veg. is vegetal, and NEB is nuclear envelope breakdown. Arrowhead marks the cytokinetic furrow. Time is in minutes:seconds. Scale bar, 10 µm **(B)** Plot of the distance between the edge of the nucleus and the center of the CC (magenta, left *y*-axis; *N* = 6 embryos, *n* = 12 cells) and centrosome separation within a CC (green, right *y-axis*; *N* = 9 embryos, *n* = 18 cells) as a function of time. The dashed line at 17 min indicates NEB. Error bars, standard error of the mean. **(C)** Brightfield and confocal maximum intensity projections of centrosomes (magenta, middle row) and microtubules (green, bottom row) during laser ablation of microtubules between the centrosomes. The yellow line demarcates the ablation site. Ant. is anterior, Veg. is vegetal. Time is in minutes:seconds. Scale bar, 30 µm. Inset scale bar, 5 µm. **(C’)** Plot of the rotation and tilt angles between the 2-cell stage CCs or spindles and the 1-cell stage spindle as a function of time in control embryos (*N* = 9 embryos, *n* = 18 cells), ablated cell (*N* = 5 embryos, *n* = 5 cells), and non-ablated cell (*N* = 5 embryos, *n* = 5 cells) of laser-ablated embryos between the centrosomes. The dashed line at 17 min indicates NEB. Error bars, standard error of the mean. **(D)** Brightfield and confocal maximum intensity projections of the DNA (magenta) and microtubules (green) during laser ablation of microtubules between the nucleus and the CC. The yellow line demarcates the ablation site. Ant. is anterior, Veg. is vegetal. Time is in minutes:seconds. Scale bar, 30 µm. Inset scale bar, 5 µm. **(D’)** Plot of the rotation and tilt angles between the 2-cell stage CCs or spindles and the 1-cell stage spindle as a function of time in control embryos (*N* = 9 embryos, *n* = 18 cells), ablated cell (*N* = 4 embryos, *n* = 4 cells), and non-ablated cell (*N* = 4 embryos, *n* = 4 cells) of laser-ablated embryos between the centrosomes and the nucleus. The dashed line at 17 min indicates NEB. Error bars, standard error of the mean.

Because centrosome separation initiates when the nucleus reaches them, we assessed whether the CCs use the nuclei to orient/power their movements as previously observed in *C. elegans* and *Drosophila* embryos (Bondaz et al., 2019; Buttrick et al., 2008; Meyerzon et al., 2009; Tanenbaum and Medema, 2010). To test this possibility, we performed periodic laser ablations every 90 s between the CC and the nucleus in one of the two cells from the time when both centrosomes were identifiable until the nuclei reached the center of the CC (Fig. 2D, movie 4). We found that these ablations impaired nucleus migration (Fig. S2C) but did not affect either CC movements or their final orientation (Fig. 2D’, S2D), as the CCs in sister cells of ablated embryos eventually became parallel and coplanar as in control embryos.

Altogether, these results show that CCs movements initiate at late anaphase of mitosis I and proceed during cytokinesis and interphase of the 2-cell stage in ascidians. These movements do not rely on mechanisms previously described in other embryos, such as *Xenopus* embryos (Wühr et al., 2010), *C. elegans* zygotes (Meyerzon et al., 2009), or *Drosophila* blastoderm embryos (Buttrick et al., 2008; Wong et al., 2024), and they are not supported by either the nuclei or antiparallel microtubule sliding between centrosomes.

### CC movements are powered by forces generated by astral microtubules

As the aforementioned mechanisms do not appear to contribute to CC movements, we considered the astral microtubule network to be the most likely source of the forces driving CC migration. We analyzed microtubule distribution at key times during CC migration and found that the network of astral microtubules grows at anaphase of the first mitosis and expands into the whole volume of the cell during interphase of the 2-cell stage before sharply retracting at prophase (Fig. 3A). To understand how network remodeling could contribute to the migration of the CCs, we analyzed the density of microtubules from the centrosome to the cell cortex. We found that in interphase (Fig. 3A’), a dense network of microtubules is formed extending from the centrosomes across the central cytoplasm (Fig. 3A’, blue shaded area), surrounded by a less dense peripheral network of microtubules near the cortex (Fig. 3A’, red shaded area). Upon mitotic entry at the 2-cell stage, this organization was lost, and the microtubule network became restricted to the pericentrosomal region, with the peripheral population markedly diminished (Fig. 3A’’). The long astral microtubules observed during interphase may be the sites of force generation to move the CCs either by cytoplasmic pulling, as demonstrated in sea urchin embryos (Minc et al., 2011; Minc and Piel, 2012), or by pulling of astral microtubules on the cell cortex previously shown in ascidian zygotes (Rosfelter et al., 2024), or both. To test the involvement of astral microtubules in the orientation of the CCs, we performed periodic laser ablations (every 90 s) of astral microtubules in one of the two sister cells (Fig. 3B, movie 5) from the time when both centrosomes in each CC were identifiable (late anaphase of the 1-cell stage) until NEB of the 2-cell stage. We found that the rotation angle of the CCs was severely affected in the ablated cell but not in the non-ablated one (Fig. 3B’), and that the sister CCs, and thus the sister spindles, at the 2-cell stage were no longer parallel (Fig. S3A – Rotation Ablated). In contrast, the tilt angle was not affected by these laser ablations (Fig. 3B’), and the CCs and spindles remained coplanar (Fig. S3A – Tilt Ablated). To confirm these observations, we interfered with astral microtubules using the drug 2-methoxyestradiol (2ME), which preferentially affects dynamic microtubules without impairing spindle formation (Kamath et al., 2006) (Fig. 3C). Incubating zygotes at anaphase of the 1-cell stage with 2ME drastically reduced the size of centrosomal asters during interphase of the 2-cell stage (Fig. 3C) and prevented CC migration, leading to strong defects in both rotation and tilt angles of the spindles at the end of 2-cell stage and resulting in a loss of coplanarity (Fig. 3C’-C’’, S3B-B’).

**Figure 3:**
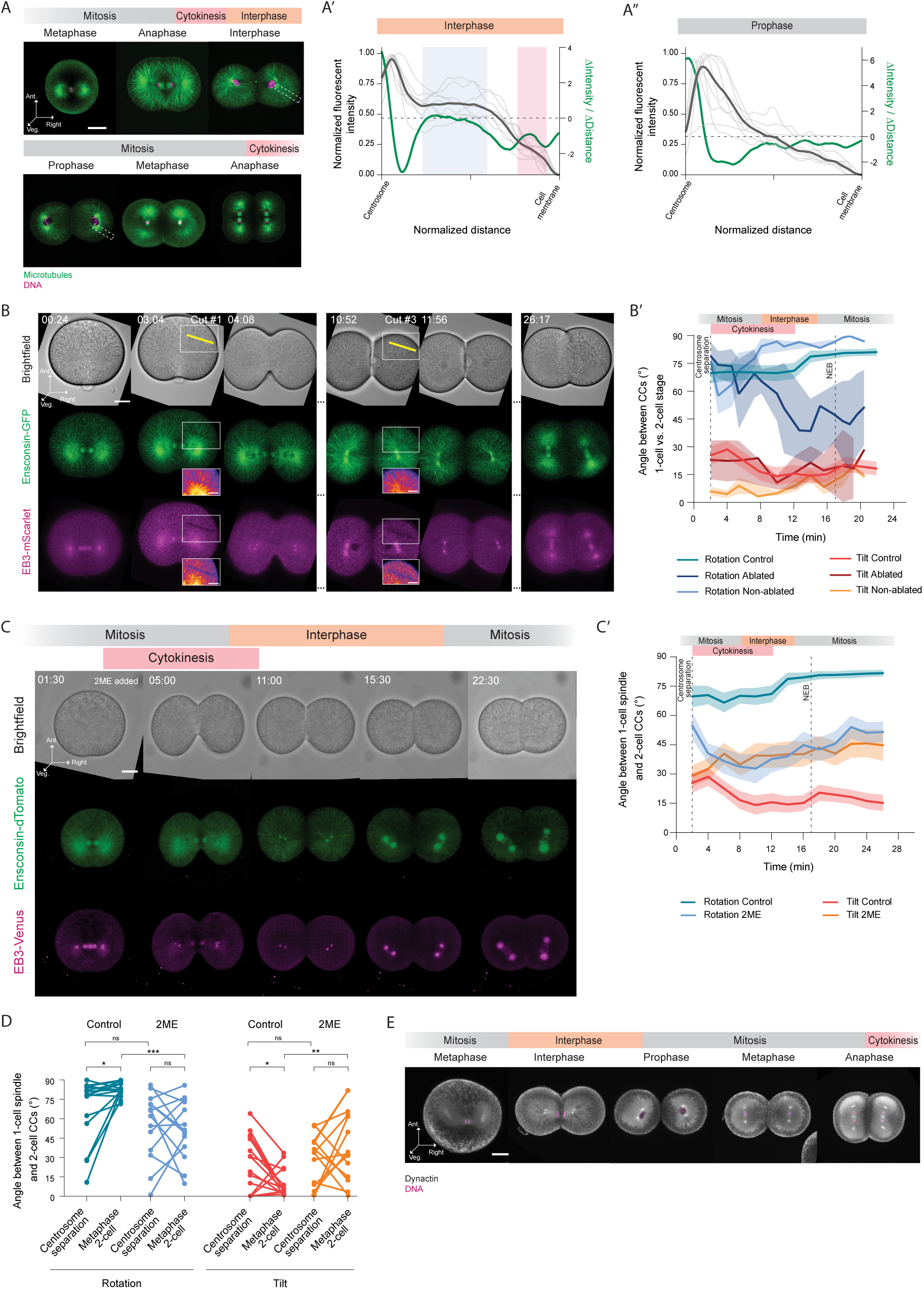
Forces along astral microtubules are essential for the orientation of CCs. **(A)** Confocal fluorescent images of a central z-slice of ascidian embryos from 1-cell stage metaphase to 2-cell stage anaphase stained for microtubules (green) and DNA (magenta). Ant. is anterior, Veg. is vegetal. The dashed area indicates the area analyzed in A’ and A’’. Scale bar, 40 µm. **(A’**, **A’’)** Fluorescent intensity profiles show the distribution of microtubules from the centrosome to the cell membrane in embryos in interphase (left *y* axis; *N* = 9 embryos) and in prophase (left *y* axis; *N* = 8 embryos). The green line (right *y* axis) represents the change in the average intensity profile over the distance. **(B)** Brightfield and confocal maximum intensity projections of microtubules (green, middle row) and centrosomes (magenta, bottom row) during laser ablation of astral microtubules. The yellow line demarcates the ablation site. Ant. is anterior, Veg. is vegetal. Time is in minutes:seconds. Scale bar, 30 µm. Inset scale bar, 20 µm. **(B’)** Plot of the rotation and tilt angles between the 2-cell stage CCs or spindles and the 1-cell stage spindle as a function of time in control embryos (*N* = 9 embryos, *n* = 18 cells), ablated cell (*N* = 4 embryos, *n* = 4 cells), and non-ablated cell (*N* = 4 embryos, *n* = 4 cells) of laser-ablated embryos. The dashed line at 17 min indicates NEB. Error bars, standard error of the mean. **(C)** Brightfield and confocal maximum intensity projections of microtubules (green, middle row) and centrosomes (magenta, bottom row) in 2ME-treated embryos. Ant. is anterior, Veg. is vegetal. Time is in minutes:seconds. Scale bar, 30 µm. **(C’)** Plot of the rotation and tilt angles between the 2-cell stage CCs or spindles and the 1-cell stage spindle as a function of time in control embryos (*N* = 9 embryos, *n* = 18 cells) and 2ME-treated embryos (*N* = 7 embryos, *n* = 14 cells). The dashed line at 17 min indicates NEB. Error bars, standard error of the mean. **(C’’)** Plot of the initial (Centrosome separation) and final (Metaphase 2-cell) rotation (left) and tilt (right) angles between the 2-cell stage CCs or spindles at 2-cell stage with respect to the 1-cell stage spindle in control embryos (*N* = 9 embryos, *n* = 18 cells; rotation: paired Wilcoxon test, *P=0.0131; tilt: paired Wilcoxon test, *P=0.0250) and 2ME-treated embryos (*N* = 7 embryos, *n* = 14 cells; rotation and tilt: paired t test, ns). Control vs. 2ME rotation angle: Kruskal-Wallis test (***P=0.0001) with Dunn’s multiple comparisons test (ns, ***P=0.0001). Control vs. 2ME tilt angle: Kruskal-Wallis test (*P=0.0101) with Dunn’s multiple comparisons test (ns, **P<0.0025). ns, not significant. **(D)** Confocal fluorescent images of a central z-slice of ascidian embryos from 1-cell stage metaphase to 2-cell stage anaphase stained for dynactin (grey) and DNA (magenta). Ant. is anterior, Veg. is vegetal. Scale bar, 30 µm.

Altogether, these results demonstrate that forces exerted on astral microtubules are responsible for the proper migration and orientation of the CCs, and thus the spindles, in 2-cell stage ascidian embryos. In other systems, these forces are generated by dynein motors either anchored at the plasma membrane, which exert tension on astral microtubules as they move toward their minus ends (Anjur-Dietrich et al., 2024; Kiyomitsu, 2019; Laan et al., 2012; Omer et al., 2018), or bound to intracellular organelles such as the ER or endosomes (Cheng and Ferrell, 2019; Shamipour et al., 2021; Woźniak et al., 2009). To explore the contribution of these mechanisms to the shaping of planar ascidian 4-cell stage embryos, we sought to localize intracellular dynein by immunocytochemistry against the dynein co-factor, dynactin. Using ascidian-specific anti-dynactin antibody (Fig. 3D, S3C), we found that dynactin is enriched both in the pericentrosomal cytoplasm and at the cell cortex. These two pools of dynactin closely correlate with the dense central and the peripheral microtubule populations, respectively, observed in interphase cells (Fig. 3A, A’).

Together, these findings suggest that dynein–dynactin complexes are enriched on locations supporting a role for both cytoplasmic and cortical pulling in the positioning of CCs and spindles. We then sought to discriminate the role of each mechanism in the positioning of the CCs.

### An anisotropic ER domain may drive CC movements in the ascidian embryo

Recent work indicates that the cytoplasm of ascidian embryos is a dense mixture of yolk granules and organelles (Stoev et al., 2025) and that the interaction between cytoskeletal structures and cytoplasmic components generates forces that shape the embryo and drive cytoplasmic organizations (Caballero-Mancebo et al., 2024). Moreover, previous studies have shown that the ER is transported by dynein motors and is able to generate pulling forces on astral microtubules (Lane and Allan, 1999; Mukherjee et al., 2020; Woźniak et al., 2009). In ascidian embryos, the ER and yolk granules segregate in the cytoplasm with the ER accumulating around the sperm aster in the zygote, while yolk vesicles are excluded from the pericentrosomal region (Rosfelter et al., 2024). We thus asked if the ER could serve as the guiding cue for the CCs to orient. We first analyzed the distribution of ER over time and found that the mitotic spindle at the 1-cell stage and the CCs during the 2-cell stage were surrounded by an accumulation of ER (Fig. 4A, S4A, movie 6), closely resembling both the pericentrosomal dense astral microtubule population and the localization of dynactin (Fig. 3A, D, and S4A). Segmentation and morphometric analysis of these ER domains revealed that, unlike the roughly spherical shape of the cell (0.85 ± 0.05 sphericity during interphase), the ER forms a cytoplasmic domain with an ellipsoid shape (0.62 ± 0.04 sphericity) and a clearly defined long axis (Fig. 4B).

**Figure 4:**
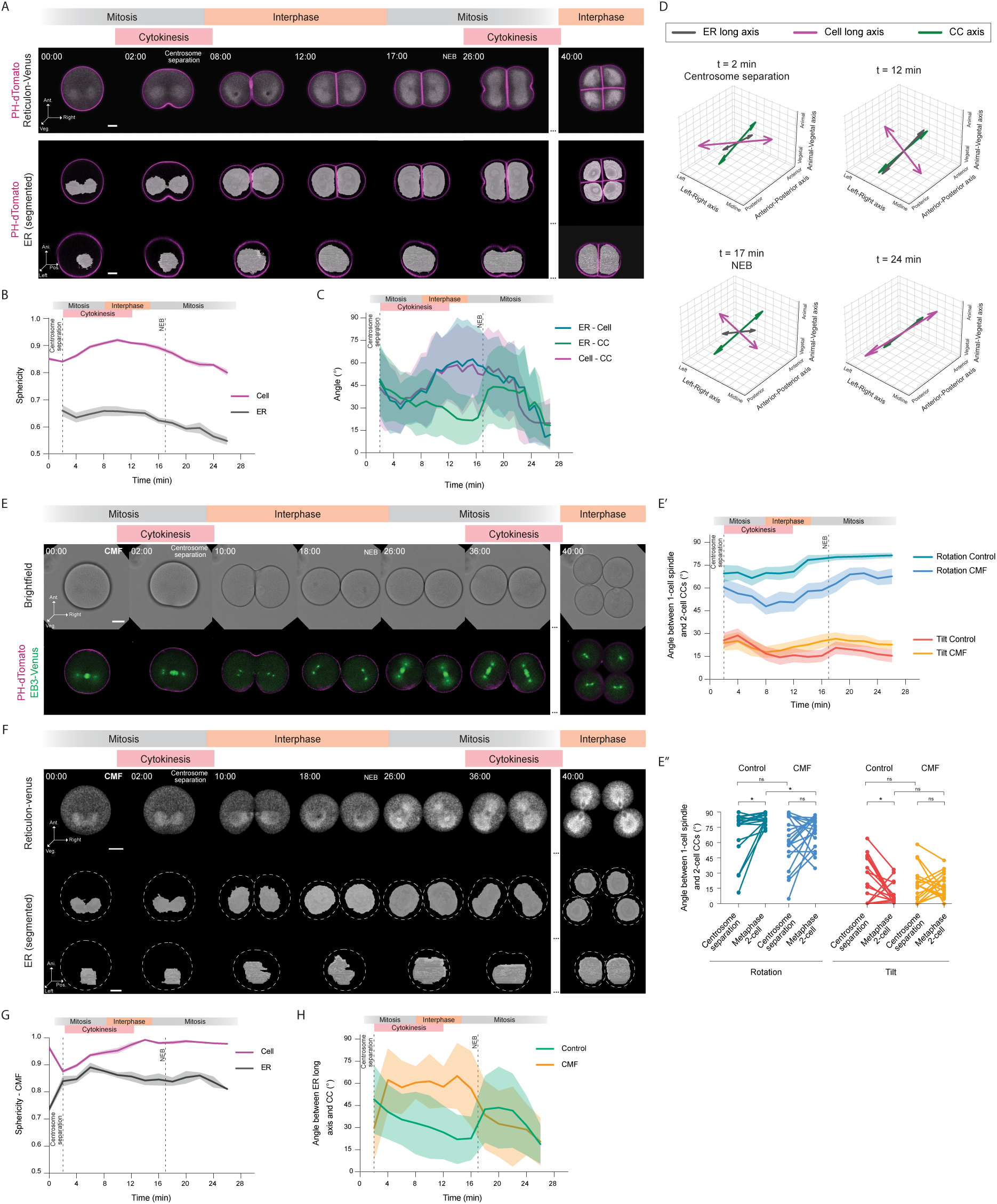
Cytoplasmic pulling controls rotation of the CCs. **(A)** Confocal fluorescent images of a central z-slice of the plasma membrane (magenta) and the endoplasmic reticulum (grey) in an ascidian embryo from 1-cell stage metaphase to 4-cell stage interphase. Middle and bottom rows show a top and side view, respectively, of the membrane (magenta) and the segmented ER (grey). Ant. is anterior, Veg. is vegetal, Ani. is animal, Post. is posterior, and NEB is nuclear envelope breakdown. Time is in minutes:seconds. Scale bar, 30 µm. **(B)** Plot of the sphericity of the cell (magenta; *N* = 14 embryos) and the ER (grey; *N* = 12 embryos) as a function of time in control embryos. The dashed line at 17 min indicates NEB. Error bars, standard error of the mean. **(C)** Plot of the angle between the long axes of the ER and the cell (blue), the ER and the CCs (green), and the cell and the CCs (red) as a function of time in control embryos. The dashed line at 17 min indicates NEB. Error bars, standard deviation (s.d.). Mean and s.d. were estimated by bootstrap method (5000 iterations). **(D)** 3D plots of the average long axes of the cell (magenta), the ER (grey), and the CCs (green) at 2, 12, 17, and 24 minutes of the left cells of ascidian embryos after metaphase of the 1-cell stage. NEB is nuclear envelope breakdown. For visualization purposes, the length of the long axes of the cell and the CCs is doubled. **(E)** Brightfield, confocal fluorescent images of a central z-slice of the plasma membrane (magenta), and maximum intensity projections of centrosomes (green) in an ascidian embryo in CMF seawater from 1-cell stage metaphase to 4-cell stage interphase. Ant. is anterior, Veg. is vegetal, Ani. is animal, Post. is posterior, and NEB is nuclear envelope breakdown. Time is in minutes:seconds. Scale bar, 40 µm. **(E’)** Plot of the rotation and tilt angles between the 2-cell stage CCs or spindles and the 1-cell stage spindle as a function of time in control embryos (*N* = 9 embryos, *n* = 18 cells) and CMF-treated embryos (*N* = 10 embryos, *n* = 20 cells). The dashed line at 17 min indicates NEB. Error bars, standard error of the mean. **(E’’)** Plot of the initial (Centrosome separation) and final (Metaphase 2-cell) rotation (left) and tilt (right) angles between the 2-cell stage CCs or spindles and the 1-cell stage spindle in control embryos (*N* = 9 embryos, *n* = 18 cells; rotation: paired Wilcoxon test, *P=0.0131; tilt: paired Wilcoxon test, *P=0.0250) and CMF-treated embryos (*N* = 10 embryos, *n* = 20 cells; rotation and tilt: paired t test, ns). Control vs. CMF rotation angle: Kruskal-Wallis test (*P=0.0194) with Dunn’s multiple comparisons test (ns, *P=0.0276). Control vs. CMF tilt angle: Kruskal-Wallis test (ns) with Dunn’s multiple comparisons test (ns). ns, not significant. **(F)** Confocal fluorescent images of the endoplasmic reticulum (top row) and its segmentation (middle row is the top view and bottom row is the side view) in an ascidian embryo from 1-cell stage metaphase to 4-cell stage interphase in CMF seawater. Ant. is anterior, Veg. is vegetal, Ani. is animal, Post. is posterior, and NEB is nuclear envelope breakdown. Time is in minutes:seconds. Scale bar, 30 µm. **(G)** Plot of the sphericity of the cell (magenta; *N* = 4 embryos) and the ER (grey; *N* = 10 embryos) as a function of time in CMF-treated embryos. The dashed line at 17 min indicates NEB. Error bars, standard error of the mean. **(H)** Plot of the angle between the long axis of the ER and the CCs or spindles as a function of time in control embryos (green, taken from C) and in CMF-treated embryos (orange). The dashed line at 17 min indicates NEB. Error bars, standard deviation.

The observed enrichment of dynactin and ER over time around the CCs points to the possibility of the CCs leveraging the anisotropic shape of the ER domain to align with its long axis, as predicted by length-dependent pulling on microtubules to position asters and spindles (Kimura and Kimura, 2011b), even in the absence of a geometrically defined long axis of the cell. To address this hypothesis, we quantified the relative alignment between the long axes of the ER domain and the cell, and the CCs using bootstrap statistics (Fig. 4C). We found that initially, the long axes of the cell, ER, and the CCs are not aligned to each other (Fig. 4C – 2 min, 4D – 2 min), displaying random orientations, which result in an average angle of ∼45°. As cytokinesis and interphase progress, the angle between the long axes of ER and the CCs gradually decreases, indicating progressive alignment between them before becoming parallel at the end of interphase (Fig. 4C – 12 min, green; 4D – 12 min, green and grey). In contrast, neither the long axis of the ER domain nor that of the CCs aligns with the long axis of the cell over the same period (Fig. 4C), suggesting that cell shape does not provide a dominant geometrical cue for CC orientation. Interestingly, at NEB, coinciding with the shortening of astral microtubules, the ER and the CCs become momentarily misaligned (Fig. 4C – 17 min, 4D – 17 min), before realigning with the cell’s long axis during mitosis of the 2-cell stage. Together, this analysis reveals that the shape of the cell does not provide a consistent geometrical cue for CC orientation, and suggests that, instead, the ER acts as a cytoplasmic scaffold that guides CC alignment independently of cell geometry, even though the final orientation of the spindles at the 2-cell stage obeys the Hertwig rule (Dumollard et al., 2017).

Even though the shape of the 2-cell stage blastomeres remained fairly spherical, a long axis may still be sensed by microtubules to position the CCs and spindles. To further test whether the ER–CC alignment depends on external cell geometry or arises intrinsically from cytoplasmic organization, we cultured the embryos in calcium and magnesium-free seawater (CMF) to remove the contribution of cell adhesion to cell shape (Fig. 4E-F, movie 7). The blastomeres of CMF-treated embryos remained almost perfectly spherical, displaying an average sphericity of 0.96 ± 0.04 (Fig. 4G). The rotation angle between the CCs at the 2-cell stage relative to the mitotic spindle of the 1-cell stage, however, was significantly altered, whereas the tilt angle remained unaffected (Fig. 4E’–E’’). The defective rotation of the CCs at the end of the 2-cell stage prevented the mitotic spindles from being parallel at mitosis (Fig. S4B-C) and affected the characteristic square geometry of 4-cell stage embryos. Planarity of the embryos was, however, preserved in CMF-treated embryos (Fig. S4B-C). We then analyzed the shape of the ER domain around the CCs in CMF-treated embryos (Fig. 4F) and found that it was less ellipsoidal than in control embryos - average sphericity of 0.91 ± 0.04 (Fig. 4G) - supporting the idea that the ER can mirror the cell shape anisotropy. Moreover, we found that the angle between the long axis of the ER and the CCs was consistently higher than in control embryos, indicating reduced alignment between them (Fig. 4H). This suggests that cell geometry primarily impacts the progressive alignment of the CCs through rotation by cytoplasmic-generated forces, whereas the tilt angle does not seem to be influenced by cell shape.

### Anisotropic cortical pulling forces maintain the CCs coplanar during CCs migration

We next asked if cortical pulling forces control the posterior tilt along the anterior-posterior axis (Fig. 1A, D). To that end, we first monitored the evolution of cortical pulling forces during CC migration by imaging plasma membrane invaginations towards the centrosomes after cortical softening with 1 µg/ml Latrunculin B (Fig. 5A-B, movie 8), an indication of cortical pulling sites (Godard et al., 2021; Rosfelter et al., 2024). We found that plasma membrane invaginations appear at anaphase of the 1-cell stage and that their frequency increases during interphase of the 2-cell stage before sharply decreasing at prophase and metaphase of the 2-cell stage mitosis (Fig. 5B’), similarly to what has been found in the zygote (Rosfelter et al., 2024). These observations show a good temporal correlation between the appearance of the network of long astral microtubules and the occurrence of cortical pulling, supporting the idea that cortical pulling may also be responsible for CC movements. We then looked at the spatial distribution of cortical pulling sites and found that membrane invaginations were more frequent at the posterior pole of the embryo compared to the anterior pole (Fig. S5A). This difference was consistent throughout the migration of the CCs and persisted until anaphase at the 2-cell stage, supporting our hypothesis that asymmetrical cortical pulling forces may play a central role in regulating the tilt angle and coplanarity of CCs, thereby controlling the planar architecture of the 4-cell stage embryo. Surprisingly, analysis of the CCs orientation in Latrunculin B-treated embryos (Fig. S5B) showed that neither rotation nor tilt angles are affected by drastic softening of the embryo cortex (Fig. S5B’) and that the spindles at mitosis of the putative 2-cell stage remain parallel and coplanar (Fig. S5B’’). This is consistent with previous reports that reducing actomyosin contractility has minimal impact on sperm aster positioning (Rosfelter et al., 2024) or early spindle orientation (Berends et al., 2013), and that cortical pulling can persist when actin integrity is compromised (Redemann et al., 2010), while excessive cortical tension can suppress pulling (Kelkar et al., 2022). Thus, the question remains as to whether cortical pulling can regulate CCs movements.

**Figure 5:**
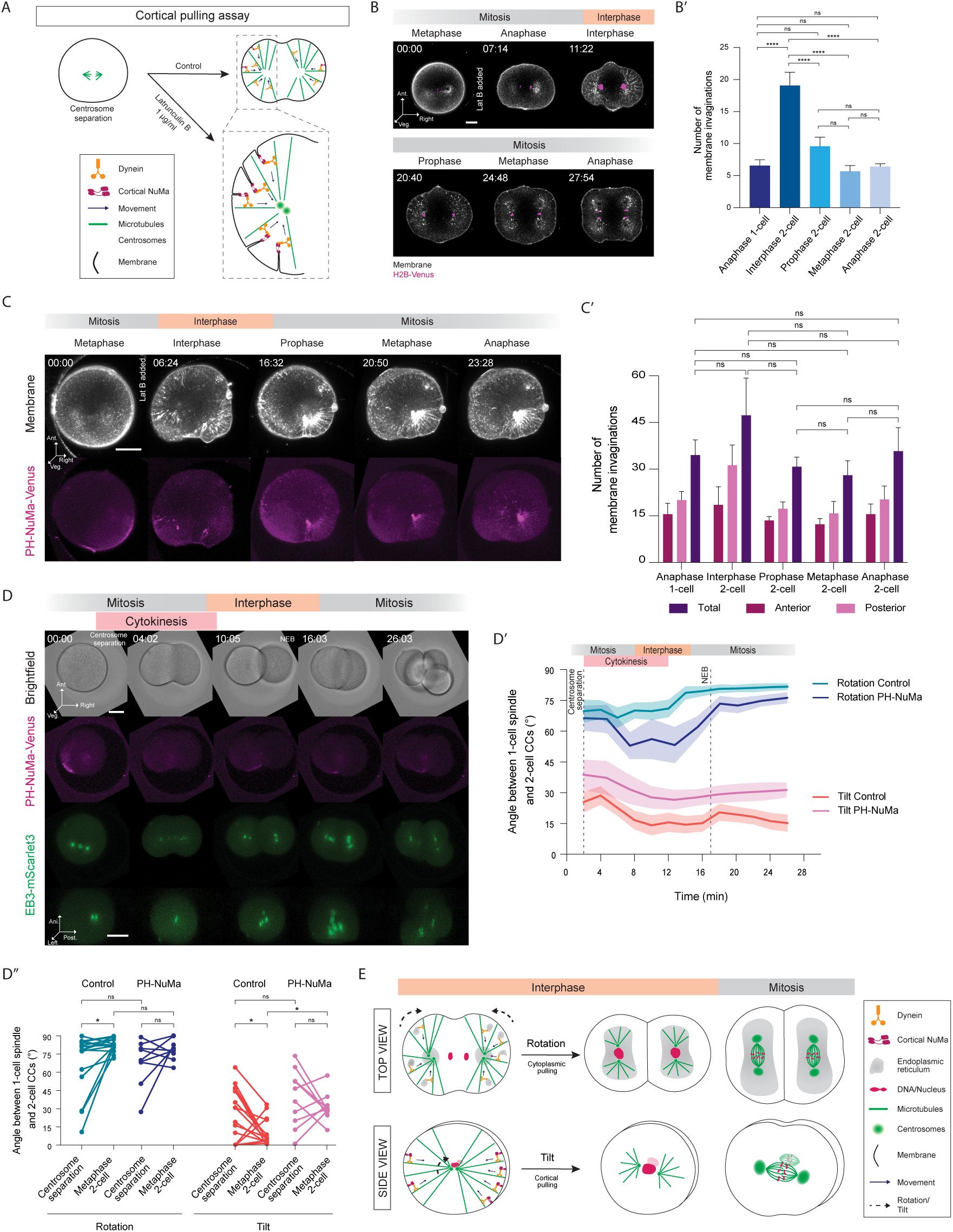
Tilt along the anterior-posterior axis is controlled by cortical pulling forces. **(A)** Schematic representation of the cortical pulling assay. **(B)** Confocal maximum intensity projections over 6 central z-slices of the plasma membrane (grey) and DNA (magenta) in an ascidian embryo from 1-cell stage metaphase to 2-cell stage anaphase, treated with 1µg/ml Latrunculin B from 1-cell stage anaphase. Ant. is anterior, Veg. is vegetal. Time is in minutes:seconds. Scale bar, 30 µm. **(B’)** Bar plot of the average number of membrane invaginations as a function of the cell cycle stage. (*N* = 13 embryos). One-way ANOVA (****P<0.0001) and Tukey’s multiple comparisons tests (****P<0.0001; ns - not significant). Error bars, standard error of the mean. **(C)** Confocal maximum intensity projections over 6 central z-slices of PH-NuMA (magenta) and membrane (grey) in an ascidian embryo expressing PH-NuMA and treated with 1µg/ml Latrunculin B from 1-cell stage anaphase. Scale bar, 40µm. **(C’)** Bar plot of the average number of membrane invaginations (anterior, posterior, and total) as a function of the cell cycle stage in embryos expressing PH-NuMA and treated with 1µg/ml Latrunculin B. (*N* = 4 embryos). Kruskal-Wallis test and Dunn’s multiple comparisons test. Error bars, standard error of the mean. ns, not significant. **(D)** Brightfield and confocal maximum intensity projections of PH-NuMA (magenta) and centrosomes (green) in embryos expressing PH-NuMA. Ant. is anterior, Veg. is vegetal, Ani. is animal, Post. Is posterior. Time is in minutes:seconds. Scale bar, 40 µm. **(D’)** Plot of the rotation and tilt angles between the 2-cell stage CCs or spindles and the 1-cell stage spindle as a function of time in control embryos (*N* = 9 embryos, *n* = 18 cells) and embryos expressing PH-NuMA (*N* = 5 embryos, *n* = 10 cells). The dashed line at 17 min indicates NEB. Error bars, standard error of the mean. **(D’’)** Plot of the initial (Centrosome separation) and final (Metaphase 2-cell) rotation (left) and tilt (right) angles between the 2-cell stage CCs or spindles and the 1-cell stage spindle in control embryos (*N* = 9 embryos, *n* = 18 cells; rotation: paired Wilcoxon test, *P=0.0131; tilt: paired Wilcoxon test, *P=0.0250) and embryos expressing PH-NuMA (*N* = 7 embryos, *n* = 14 cells; rotation and tilt: paired t test, ns). Control vs. PH-NuMA rotation angle: Kruskal-Wallis (*P=0.0201) with Dunn’s multiple comparisons test (ns). Control vs. PH-NuMA tilt angle: Kruskal-Wallis (**P=0.0073) with Dunn’s multiple comparisons test (ns, *P=0.0169). ns, not significant. **(E)** Model of the mechanisms controlling CC positioning during the 2-cell stage: cytoplasmic dynein-dependent pulling along astral microtubules controls the 90° rotation of the CCs with respect to the 1-cell stage spindle. Dynein motors on the ER surface move towards the CCs, thus forming the pericentromeric ER domains. These domains act as a proxy for cell shape, dictating the rotation of the CCs during interphase to generate parallel spindles at the 2-cell stage. Concomitantly, cortical dynein generates cortical pulling forces at the end of astral microtubules that control the tilt of the CCs, ensuring the coplanarity of sister CCs and the planar geometry of the 4-cell stage ascidian embryo.

The observation of the posterior bias in cortical pulling forces and its correlation with spindle tilt led us to hypothesize that increasing cortical pulling would affect CC orientation (Middelkoop et al., 2024) and perturb the coplanarity of the 4-cell stage embryos. In animal cells, cortical pulling forces are mediated by dynein–NuMA–LGN–Gαi complexes anchored to the plasma membrane (Anjur-Dietrich et al., 2024). In this complex, the heterotrimeric G-protein subunit Gαi recruits LGN to the cortex, which in turn binds NuMA that acts as a scaffold linking the cortical complex to astral microtubules, thereby allowing dynein motors to generate pulling forces as they move toward the microtubule minus ends. To test whether increasing cortical pulling alters CCs positioning and embryo geometry, we overexpressed NuMA and targeted it to the cortex by fusing it to a pleckstrin homology (PH) domain, which promotes membrane localization (Dumollard et al., 2017). In embryos expressing PH-NuMA, the fusion protein predominantly accumulated at the posterior cortex (Fig. 5C, bottom row, 0 min; movie 9), consistent with the accumulation of plasma membrane in the posterior part of the embryo (Carroll et al., 2003). The cortical pulling assay in these embryos confirmed a global increase in the number of plasma membrane invaginations (Fig. 5C’, Total) compared to control embryos (Fig. 5B’), with a higher frequency of posterior pulling sites, confirming that cortical pulling was strengthened and correlated with the accumulation of PH-NuMA.

Analysis of the orientation of CCs in embryos overexpressing cortical NuMA showed an increase in the tilt angle at the 2-cell stage relative to the spindle at the 1-cell stage, while rotation remained unaffected (Fig. 5D-D’’, movie 10). The tilt angle was also increased between sister CCs at the 2-cell stage (Fig. S5C, S5D - Tilt), which are not coplanar anymore, resulting in embryos adopting a more tetrahedral configuration (Fig. 5C, 26 min), in contrast to the planar geometry observed in controls. Altogether, these results suggest a model in which proper CC orientation, and thus spindle, determines the planar 4-cell embryo geometry through a finely tuned balance between cortical and cytoplasmic dynein-mediated pulling forces acting on astral microtubules in a cell cycle-dependent manner (Fig. 5E).

## DISCUSSION

Our study uncovers the mechanisms that drive the dynamic migration of centrosomal complexes (CCs) to achieve proper spindle orientation in early ascidian embryos and identifies a previously underappreciated balance between cortical and cytoplasmic forces as a key determinant of embryo geometry. Through quantitative live imaging and biophysical manipulations, we reveal that ascidian embryos rely on a finely tuned interplay between dynein-dependent forces acting from the cell cortex and cytoplasmic forces that are most likely transmitted through the ER. We demonstrate that the CCs of the two cells initially form at varying angles relative to the position of the first mitotic spindle and progressively achieve parallel alignment during interphase of the 2-cell stage. Interestingly, a second cue determined by cortical pulling forces controls the coplanarity of the CCs, ensuring the planar arrangement of the blastomeres at the 4-cell stage.

Previous studies in *C. elegans* zygotes have revealed the major role of cortical pulling forces in powering centrosome separation, which is then buffered by cytoplasmic pulling forces acting on the nuclear envelope (De Simone et al., 2016). However, centrosomes of enucleated 2-cell stage *C. elegans* embryos can separate normally without interacting with a nucleus (Fujii et al., 2023), supporting the primary role of cortical pulling forces in centrosome separation. In large cells such as *Xenopus* and zebrafish eggs, however, astral microtubules cannot extend completely to the cell cortex, rendering cortical pulling inefficient for centrosome separation and orientation (Pierre et al., 2016; Wühr et al., 2010). In these cells, length-dependent cytoplasmic pulling forces are responsible for moving and orienting centrosomes, relying on spatial cues arising from the anisotropic shape of asters sensitive to both cell shape and cytoplasmic stratification (Pierre et al., 2016). In *Xenopus* embryos, final positioning of the centrosomes is achieved before the nuclear envelope reforms, during anaphase of mitosis I, via kinesin Eg5-mediated antiparallel sliding of microtubules, which separates centrosomes (Wühr et al., 2010). In *C. elegans* embryos, Eg5 activity also cooperates with cytoplasmic pulling for centrosome separation (Bondaz et al., 2019). Our laser ablation experiments show that, in ascidian embryos, neither antiparallel forces between centrosomes nor the link with the nucleus are required for the alignment of the CCs at the 2-cell stage. This finding demonstrates that ascidian embryos employ a distinct mechanism for spindle orientation that is largely independent of the nuclear or centrosomal cues that are crucial in *Xenopus* and *C. elegans*.

Ascidian embryos are of an intermediate cell size between small nematode embryos, where cortical forces dominate, and the large *Xenopus* and zebrafish embryos, where cytoplasmic forces dictate spindle orientation. Our study reveals that the final orientation of CCs and spindles depends on pulling forces on astral microtubules from both the cortex and the cytoplasm and that their balance dictates the final shape of the 4-cell stage embryo. We show that the final orientation of the CCs is achieved progressively during cytokinesis of mitosis I and interphase of the 2-cell stage, and remains unchanged during subsequent mitosis. Concomitant with this alignment, the ER accumulates progressively around the CCs, suggesting that it is pulled towards the minus ends of microtubules. This observation, together with the fact that the microtubule network is denser in areas filled with ER and devoid of yolk platelets, leads us to propose a model in which the ER is the cytoplasmic cargo supporting length-dependent microtubule pulling (Mukherjee et al., 2020; Xie et al., 2022). Microtubule exclusion from yolk-dense regions has been found in *Xenopus* to be responsible for the geometric control of cell divisions in early embryos (Pierre et al., 2016; Wühr et al., 2010). The spatial organization of the ER provides experimental evidence that the information from cell shape is robustly transmitted to the CCs by the ER domain and powered by length-dependent pulling on the ER along astral microtubules. Therefore, a stratified cytoplasm can shape where and how intracellular forces are generated, which in turn influences spindle behavior (Pierre et al., 2016) and provides early embryos with a robust mechanism to enforce the Hertwig rule.

Our quantifications also show that the CCs in the two blastomeres remain coplanar to each other while they rotate to achieve the final alignment. We find that the tilt angle between the CCs and the mother spindle converges to 15° (and not 0°), similar to the tilt angle found in another ascidian species at the 4- to 8-cell stage cell division (Godard et al., 2021; Negishi et al., 2007). Ectopically inducing more cortical pulling increased the tilt angle difference between the two CCs and prevented 4-cell stage embryos from being planar. Moreover, the persistence of coplanar, but not square, divisions in CMF seawater-treated embryos further implies that cortical inputs primarily refine coplanarity, while ER-based cytoplasmic forces provide the dominant contribution to CC rotation and global spindle alignment.

Our findings expand the repertoire of known mechanisms by which cells can achieve precise spatial organization during development and highlight the evolutionary diversity of strategies employed by different organisms. Proper spindle orientation is not only critical for ensuring correct cleavage patterns and cell fate specification but also for the establishment of tissue architecture and organogenesis. The mechanisms uncovered in ascidian embryos may thus inform our understanding of how similar processes are regulated in other organisms, and may provide insights into the evolution of developmental strategies across metazoans.

## MATERIALS AND METHODS

### Animal Collection and Maintenance and Embryo Handling

Adult *Phallusia mammillata* were collected in Sète (France) and kept at 16 °C in the aquaria of the Centre de Resources Biologiques (CRB) of the Institut de la Mer de Villefranche (IMEV), which is an EMBRC-France certified service (https://www.embrc-france.fr/fr/nos-services/fourniture-de-ressources-biologiques/organismes-modeles/ascidie-phallusia-mammillata). All experiments were performed at 18-19 °C. Oocytes and sperm were obtained after animal dissection and used on the day of collection. Oocytes were chemically dechorionated in a 0.1 % trypsin solution (Merck, T8003) in micro-filtered natural sea water (MFSW; Nalgene 0.2 µm filters, cat. no. Z370584) for 90 min under gentle stirring and kept in MFSW until fertilization with 6 µl sperm that was activated by resuspension in 1 ml of MFSW at pH 9.2.

Embryos were cultured either in MFSW or in Ca^2+^ and Mg^2+^-free (CMF) seawater supplemented with 1 mM EDTA (Sardet et al., 2011).

### Cloning of Expression Constructs

To generate the pSPE3-EB3-mScarlet plasmid, the EB3 coding sequence was amplified from pRN3-EB3-3xVenus vector using the following primers with EcoRI and XhoI sites, respectively: 5’-GAATTCAAAATGGCCGTCAATGTGTACTCCACAT-3’ and 5’-CTCGAGGTACTCGTCCTGGTCTTCTTGTTGA-3’. The mScarlet coding sequence (including stop codon) was amplified from the plasmid pSPE3-iMyo-mScarlet (Caballero-Mancebo et al., 2024) using the following primers with XhoI and NotI sites, respectively: 5’-CTCGAGATGGTGAGCAAGGGCGAGGCAGTG-3’ and 5’-GCGGCCGCCTACTTGTACAGCTCGTCCATGCC-3’. To generate the pSPE3-EB3-mScarlet plasmid, the PCR products were ligated into a pSPE3-Rfa destination vector (Roure et al., 2007) that had been previously digested with EcoRI and NotI.

To generate the pSPE3-Reticulon-Venus plasmid, the *Phallusia mammillata* reticulon coding sequence (GenBank CAB3265822.1) was amplified from a cDNA library using the following primers: 5’-CACCATGGAGTATGAAACGAGCCC-3’ and 5’-CTCTGCTTTAGGCTTGGCACC-3’. The PCR product was used to generate an entry vector using the pDONR/TOPO (p3M-Reticulon). The entry vector was recombined with the destination vector pSPE3-Rfa-Venus (Roure et al., 2007).

To generate the pSPE3-PH-NuMA-Venus plasmid, the *Phallusia mammillata* NuMA coding sequence (GenBank CAB3265828.1) was amplified from a cDNA library using hybridizing primers: 5’-GAGCTCAACATCATGGATGATGACGAAA-3’ and 5’-GTCTGAATCCAAAAAAGAAGGGTGGGCG-3’. The PH coding sequence was amplified from pSPE3-PH-dTomato plasmid (Prodon et al., 2010) using the following hybridizing primers: 5’-AAGCAGGCTCCGCGGCCGCCCCCTTCA-3’ and 5’-GAGCTCAACATCATGGATGATGACGAAA-3’. After hybridization, the full PH-NuMA coding sequence was amplified using the following primers: 5’-ACAAGTTTGTACAAAAAAGCAGGCTCCGCGGCCGCCCCCTTCA-3’ and 5’-GTCTGAATCCAAAAAAGAAGGGTGGGCGCGCCGACCCAGCTTTCTTGTACAAAGTGGT-3’. The final PCR product was used to generate an entry vector using the pDONR/TOPO (p3M-PH-NuMA). The entry vector was recombined with the destination vector pSPE3-Rfa-Venus (Roure et al., 2007).

### mRNA Microinjections

Microinjections were performed as described(McDougall et al., 2014). Microinjection needles (Harvard Apparatus, 30-0035) were pulled with a PN-30 puller (Narishige) and mounted on a micromanipulation setup (MMO-203, Narishige) attached to an Olympus IX70 microscope. Dechorionated unfertilized eggs were placed into a glass wedge mounted onto a 400 µl Perspex mounting chamber (McDougall et al., 2014). *In vitro* mRNA synthesis was performed using the mMESSAGE mMACHINE T3 Transcription Kit (Ambion, AM1343). The following mRNAs were injected: EB3-3xVenus (Rosfelter et al., 2024), EB3-mScarlet (this study), Ensconsin-3xGFP (Costache et al., 2017), Ensconsin-dTomato (Costache et al., 2017), H2B-Venus (Prodon et al., 2010), PH-dTomato (Prodon et al., 2010), PH-NuMA-Venus (this study), and Reticulon-Venus (this study). mRNAs were injected at a concentration of 2-4 µg/µl using a high-pressure system (Narishige IM300) and incubated at 16 °C overnight before the experiment.

### Myoplasm and Plasma Membrane Labeling

Dechorionated unfertilized oocytes or zygotes were incubated in a 1 μM solution of MitoTracker™ Deep Red FM (Thermo Fisher, M22426) to label the myoplasm and/or in a 1 μM solution of CellMask™ Orange (Thermo Fisher, C10045) in MFSW for 5 min and washed in MFSW before imaging.

### Cytoskeletal Inhibitors

To soften the cortex, 1 μg/ml Latrunculin B (Merck, L5288) was added to the imaging dish 40 minutes post fertilization (right before first cell division) and left in the medium for the remainder of the experiment.

To interfere with dynamic microtubules, 1 μg/ml 2-methoxyestradiol (Merck, M6383) was added to the imaging dish 40 minutes post fertilization (right before first cell division) and left in the medium for the remainder of the experiment.

### Confocal Live Imaging

Embryos were mounted 30 minutes post fertilization in glass-bottom dishes (Cellvis, D35-10-0-N) coated with 1% low gelling temperature agarose (Merck, A2576) in MFSW. Embryos were imaged in either an inverted Leica TCS SP8 confocal microscope or an inverted Leica Stellaris confocal microscope using a HCPL APO CS2 40x/1.1 NA water immersion objective (Leica) and Hybrid detectors. Embryos were imaged *in toto* with an optical section thickness of 3 μm every 120-180 s. For membrane invagination experiments, the frame rate was reduced to 30 s, and for fixed samples, the optical section was 1 μm.

### Laser Ablation

Embryos were mounted 30 minutes post fertilization as for confocal live imaging and imaged on a Zeiss LSM980 confocal microscope equipped with a two-photon pulsed laser using a C-Aprochromat 40x/1.1 NA water immersion objective. Microtubules between centrosomes or between centrosomes and the nucleus were cut along a 10 μm line and astral microtubules were cut along 50 μm lines in 3 z planes, all of which were performed with a two-photon Coheren Axon 920-2 laser (2.5W). For ablation, the two-photon laser was used at full power, and the power received at the focal point is estimated at 2.8 ·10^12^ W/m^2^. Embryos were imaged over 50 central z-slices with an optical section thickness of 1 μm every 13 s.

### Antibody Production

The ascidian-specific anti-dynactin mouse polyclonal antibody was produced by fusing 1002 base pairs, amino acids 35 to 334, of the *Ciona intestinalis* dynactin subunit 1 protein (GenBank AP042712.1) to a 6-His tag at the C terminus and cloned into the vector pET11. Expression in bacteria was induced with IPTG, and the dynactin fusion protein was purified on a nickel column and eluted with imidazole according to the manufacturer’s instructions (Potrino Ni-NTA column, Macherey-Nagel). Immunization of mice was performed by Covalab (France), and sera was obtained after 57 days were used directly without further purification.

### Western Blot

Anti-dynactin antibody’s specificity was validated by Western blot analysis showing a single band at ∼55 kDa corresponding to dynactin (Fig. S3C).

2 and 8-cell stage embryos were collected in a small volume of MFSW and mixed with the same volume of 2x Laemmli sample buffer (100 mM Tris-HCL pH 6.8, 4% SDS, 0.2% Bromophenol Blue, 20% glycerol and 200 mM dithiothreitol). Samples were heated for 5 min at 95 °C and then stored at −20 °C. Proteins were separated by electrophoresis on SDS-polyacrylamide gels and transferred to a nitrocellulose membrane. Membranes were stained briefly with Ponceau-S, then washed with TBS-Tw (20 mM Tris Base and 150 mM NaCl plus 0.1% Tween-20), and blocked for at least 1 hour in TBS-Tw containing 5% dry milk powder. Immunoblots were then incubated in anti-dynactin primary antibody diluted 1:1000 overnight at 4 °C. Membranes were then washed three times with TBS-Tw and incubated with an anti-mouse horseradish peroxidase-conjugated secondary antibody (Jackson ImmunoResearch, 115-035-003) at 1:10,000 dilution in TBS-Tw + 5% dry milk powder for 2 hours at room temperature. After three washes in TBS-Tw, signal detection was carried out using the SuperSignal^TM^ West PLUS Pico chemiluminescent substrate (ThermoFisher Scientific, 34580), and images were acquired with a Fusion FX system (Vilber Lourmat, Collegien, France).

### Immunostaining

Embryos were fixed in 100% methanol (VWR Chemicals, 67-56-1), which had been pre-chilled at −20°C, and then stored at −20°C. Samples were rehydrated by three washes in PBS containing 0.1% Tween (PBS-Tw), then blocked in PBS-Tw containing 1% BSA for 1 hour at room temperature. Primary antibodies to tyrosinated alpha tubulin (rat anti-alpha-tubulin tyrosinated, monoclonal YL1/2, Merck, MAB1864-I) or to dynactin subunit 1 (mouse polyclonal, custom made, see above) were added at a dilution of 1/500 in blocking buffer and incubated for at least 24 hours at 4 °C. After incubation, samples were washed three times in PBS-Tw, and fluorescent anti-rat (Jackson ImmunoResearch, 112-095-003) or anti-mouse secondary antibodies (Jackson ImmunoResearch, 115-175-146) were added at a dilution of 1/200. After further incubation at room temperature for 1-2 hours, samples were washed in PBS-Tw, labelled with Hoechst 33342 (final concentration 5 µg/ml, Merck, TA9H97BAECD2) for 10 minutes, washed twice more in PBS-Tw, and mounted on glass slides in CitiFluor AF1 (Electron Microscopy Services, 17970-25). To prevent embryo compression or breakage, coverslips were elevated by application of small spacers of modelling clay at the corners.

### 3D Tracking of Centrosomes and Angle Quantification

Confocal fluorescent z-stacks of embryos injected with EB3-3xVenus and PH-dTomato were stabilized using the MultiStackReg (Thevenaz et al., 1998) plugin in Fiji (Schindelin et al., 2012). Centrosomes were then tracked using the MTrackJ (Meijering et al., 2012) plugin in Fiji, and their 3D coordinates (*Ci*_*x*_, *Ci*_*y*_, *Ci*_*z*_) were used to calculate the polar angles (rotation and tilt) with the y-axis (embryonic anterior-posterior axis) as reference.

### Centrosome separation

The 3D coordinates of each centrosome were used to calculate the distance between each centrosome within a 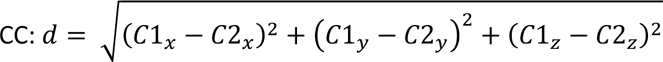

### Distance Between the Nucleus and the CC

Confocal fluorescent z-stacks of embryos injected with H2B-Venus and EB3-mScarlet were stabilized using MultiStackReg plugin in Fiji. Centrosomes were then tracked using the MTrackJ plugin in Fiji, and their 3D coordinates (*Ci_x_*, *Ci_y_*, *Ci_z_*) used to calculate the midpoint of the CC, 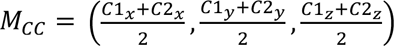. The nucleus of each cell was segmented and tracked using the Surface tool in Imaris (Bitplane). Then the distance at every time point between the nucleus (*N_x_*, *N_y_*, *N_z_*) and the CC midpoint was calculated: 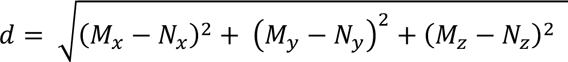. The radius of the nucleus was subtracted from the distance to reflect the time when the edge of the nucleus reaches the center of the CC.

### Microtubule Intensity Profile

The intensity profiles of microtubules were made using the line tool of 15-pixel thickness in Fiji from the centrosome to the plasma membrane. The intensity values shown were normalized to the maximum intensity value along each line. The distance between the centrosome and the plasma membrane was also normalized.

### Membrane Invagination Quantification

Confocal fluorescent z-stacks of embryos incubated in CellMask^TM^ Orange to label the plasma membrane were stabilized using MultiStackReg plugin in Fiji. 10 confocal z-slices over the equator of the embryo were sum projected. The number of membrane invaginations was counted manually by counting the invaginations present at 2-10 µm from the plasma membrane (Rosfelter et al., 2024) at the times indicated in Fig. 5B and C.

### Endoplasmic Reticulum Segmentation

Confocal fluorescent z-stacks of embryos injected with either Reticulon-Venus and PH-dTomato, Reticulon-Venus and Ensconsin-dTomato or Reticulon-Venus and EB3-mScarlet were stabilized using the MultiStackReg plugin in Fiji. For segmentation a Gaussian filter (sigma = 2) was applied and segmented using Ilastik software (Berg et al., 2019) and the pixel classification method. Segmentations were then manually corrected using Napari (Chiu et al., 2022).

### Plasma Membrane Segmentation

Confocal fluorescent z-stacks of embryos injected with PH-dTomato were stabilized using the MultiStackReg plugin in Fiji and segmented using Cellpose (Stringer et al., 2021). Segmentations were then manually corrected using Napari.

### Shape Analysis

Principal component analysis (PCA) of segmented membrane and ER timelapses was performed in order to obtain the direction of the 3 main axes using a custom-made Python script. To compute the length of the main axes (*a*, *b*, *c*), each voxel from the segmented images was projected onto each axis, and sphericity was calculated as: 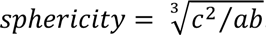.

### Statistical analysis

Statistical analysis was performed using Prism 8 (GraphPad). The statistical test used and the resulting P values are indicated in each figure legend. *N* was considered an independent experiment, and *n* denotes the number of cells; *N* and *n* are indicated in each figure. No statistical method was used to predetermine sample size. The experiments were not randomized.

## ACKNOWLEDGEMENTS

We would like to thank Hervé Turlier for fruitful discussions and the Microscopy Imaging Platform (PIM) and the animal facility (CRB) of the Institut de la Mer de Villefranche (IMEV), supported by EMBRC-France, whose French state funds are managed by the ANR within the Investments of the Future program under reference ANR-10-INBS-0. This project was funded by the Agence Nationale de la Recherche (France) grant “InvBlastula” (ANR-22-CE13-0036-01).

## AUTHORS CONTRIBUTIONS

S.C.-M., A.M., and R.D. designed the research. S.C.-M. performed the experiments and analyzed the experimental data. D.G.S. contributed to the data analysis. J.C. performed the immunostainings and antibody validations. S.C.-M. and S.B-A. performed the laser ablation experiments. L.B. created the pSPE3-EB3-mScarlet, pSPE3-Reticulon-Venus, and pSPE3-PH-NuMA-Venus constructs. S.C.-M. and R.D. wrote the manuscript.

## SUPPLEMENTAL FIGURES

**Figure S1:**
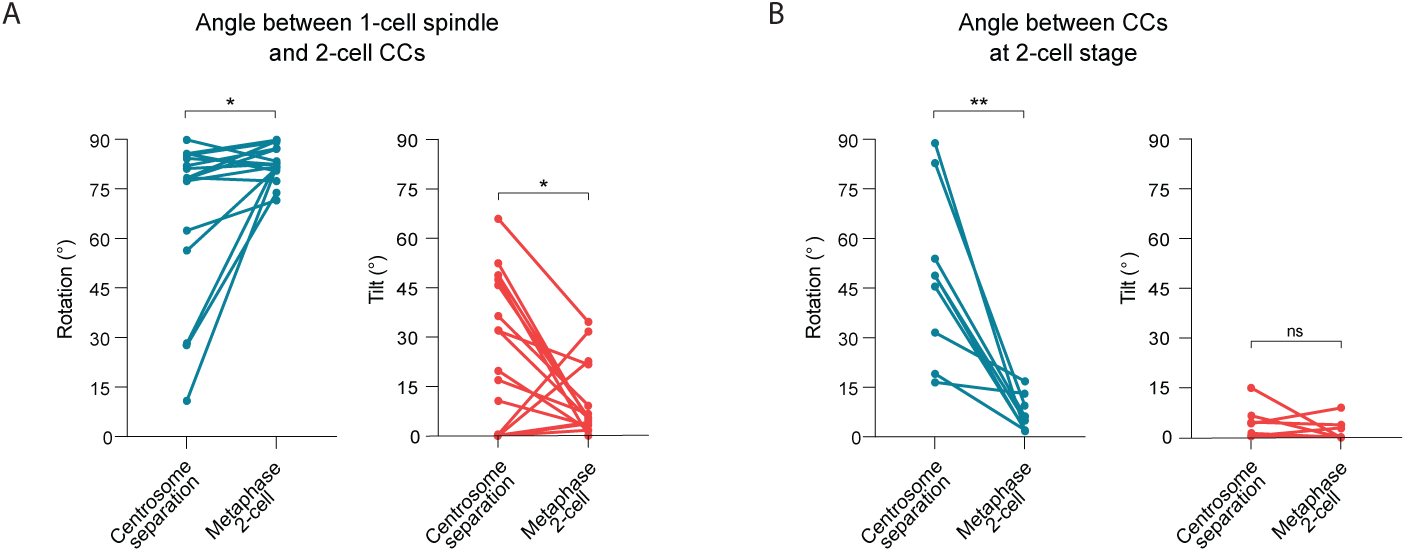
CCs of sister cells become parallel and coplanar at the 2-cell stage (relative to Fig. 1). **(A)** Plot of the initial (Centrosome separation) and final (Metaphase 2-cell) rotation (left) and tilt (right) angles between the 2-cell stage CCs or spindles and the 1-cell stage spindle (*N* = 9 embryos, *n* = 18 cells; rotation: paired Wilcoxon test, *P=0.0131; tilt: paired Wilcoxon test, *P=0.0250). **(B)** Plot of the initial (Centrosome separation) and final (Metaphase 2-cell) rotation (left) and tilt (right) angles between the left and right CCs or spindles at 2-cell stage (*N* = 9 embryos; rotation: paired t test, **P=0.0017; tilt: paired Wilcoxon test, ns – not significant).

**Figure S2:**
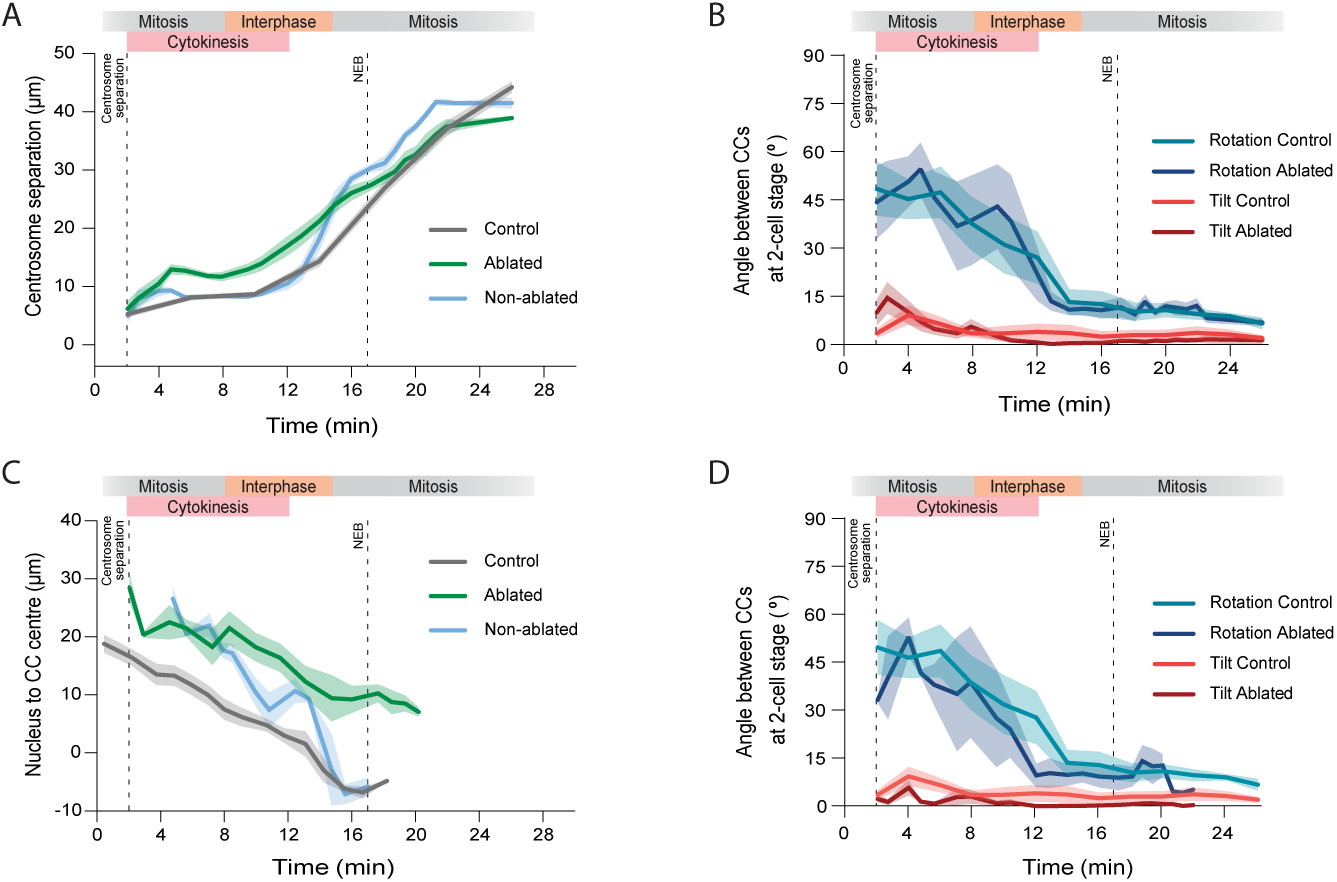
Centrosome migration and alignment in laser-ablated embryos (relative to Fig. 2). **(A)** Plot of the separation of centrosomes within a CC as a function of time in control embryos (grey; *N* = 9 embryos, *n* = 18 cells), ablated cell (green; *N* = 5 embryos, *n* = 5 cells), and non-ablated cell (blue; *N* = 5 embryos, *n* = 5 cells) of laser-ablated embryos between the centrosomes. The dashed line at 17 min indicates NEB. Error bars, standard error of the mean. **(B)** Plot of the rotation and tilt angles between the CCs or the spindles of the left and right cells at the 2-cell stage as a function of time in control (*N* = 9 embryos) and in ablated embryos between the centrosomes in one cell (*N* = 5 embryos. The dashed line at 17 min indicates NEB. Error bars, standard error of the mean. **(C)** Plot of the distance between the edge of the nucleus and the entre of the CC as a function of time in control embryos (grey; *N* = 6 embryos, *n* = 12 cells), ablated cell (green; *N* = 4 embryos, *n* = 4 cells), and non-ablated cell (blue; *N* = 4 embryos, *n* = 4 cells) of laser-ablated embryos between the centrosomes and the nucleus. The dashed line at 17 min indicates NEB. Error bars, standard error of the mean. **(D)** Plot of the rotation and tilt angles between the CCs or the spindles of the left and right cells at the 2-cell stage as a function of time in control (*N* = 9 embryos) and ablated embryos between the centrosomes and the nucleus in one cell (*N* = 4 embryos). The dashed line at 17 min indicates NEB. Error bars, standard error of the mean.

**Figure S3:**
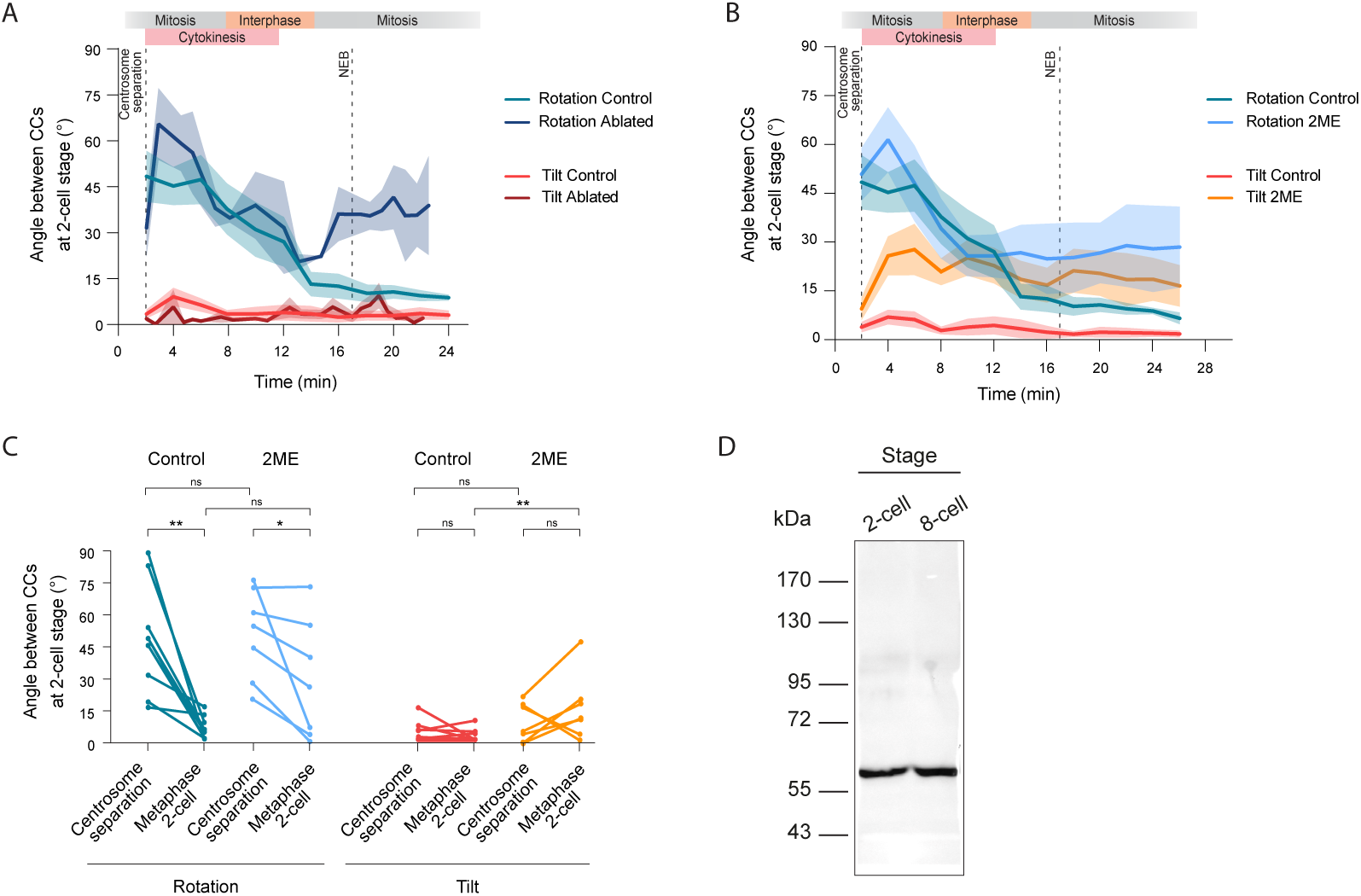
Centrosome alignment relies on forces generated along astral microtubules (relative to Fig. 3). **(A)** Plot of the rotation and tilt angles between the CCs or the spindles of the left and right cells at the 2-cell stage as a function of time in control embryos (*N* = 9 embryos), ablated cell (*N* = 4 embryos, *n* = 4 cells), and non-ablated cell (*N* = 4 embryos, *n* = 4 cells) of laser-ablated embryos. The dashed line at 17 min indicates NEB. Error bars, standard error of the mean. **(B)** Plot of the rotation and tilt angles between the CCs or the spindles of the left and right cells at the 2-cell stage as a function of time in control (*N* = 9 embryos) and 2-methoxyestradiol (2ME)-treated (*N* = 7 embryos). The dashed line at 17 min indicates NEB. Error bars, standard error of the mean. **(B’)** Plot of the initial (Centrosome separation) and final (Metaphase 2-cell) rotation (left) and tilt (right) angles between the left and right CCs or spindles at 2-cell stage in control embryos (*N* = 9 embryos; rotation: paired Wilcoxon test, *P=0.0131; tilt: paired Wilcoxon test, *P=0.0250) and embryos treated with 2ME (*N* = 7 embryos; rotation: paired t test (*P=0.0462); tilt: paired t test (ns – not significant)). Control vs. 2ME rotation angle: one-way ANOVA (***P=0.0005) with Šídák’s multiple comparisons test (ns – not significant). Control vs. 2ME tilt angle: Kruskal-Wallis test (*P=0.0186) with Dunn’s multiple comparisons test (ns – not significant, **P=0.0040). **(C)** Western blot of extracts of *Phallusia mammillata* 2- and 8-cell stage embryos probed with anti-dynactin antibody.

**Figure S4:**
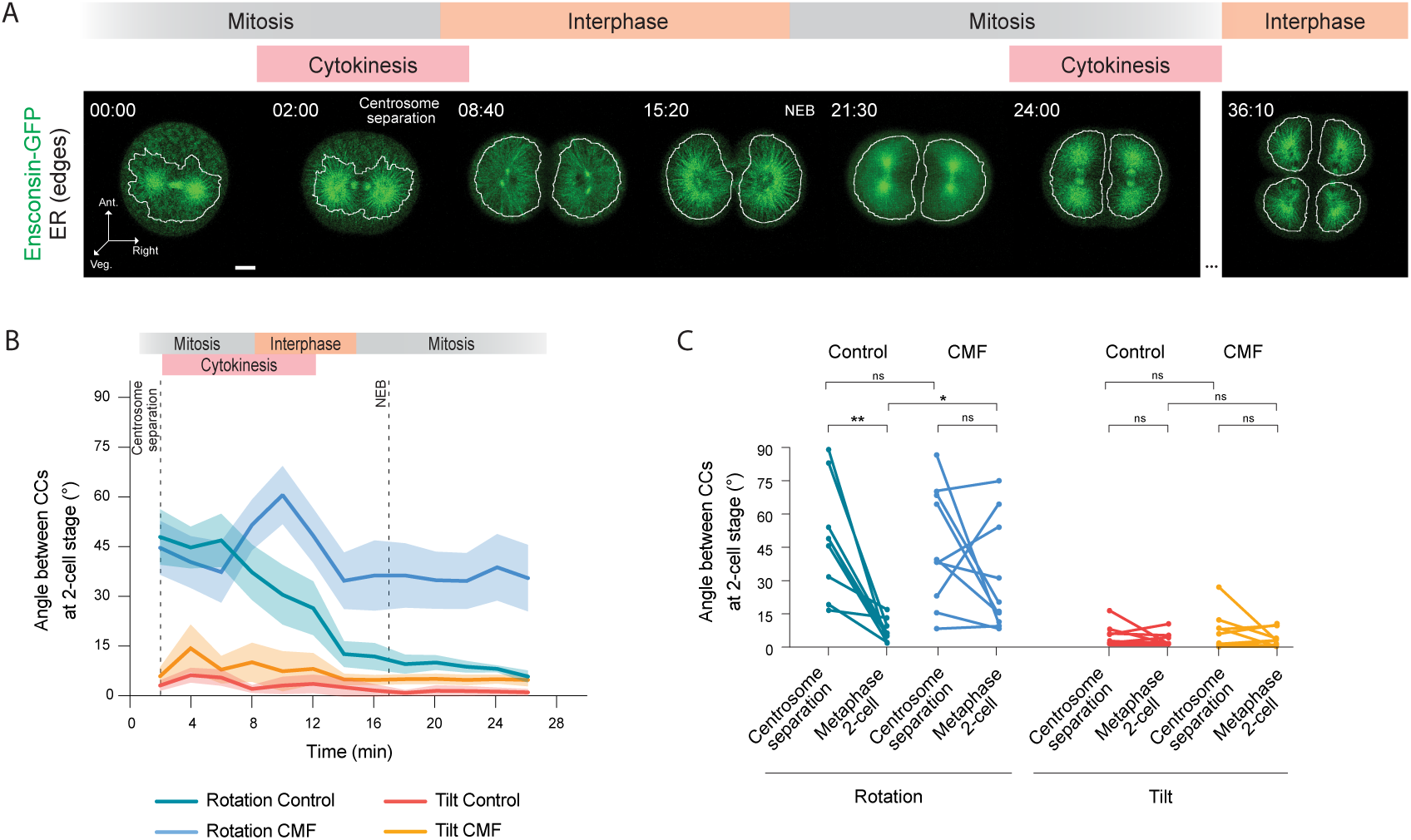
Cytoplasmic pulling controls rotation angles (relative to Fig. 4). **(A)** Maximum intensity projections of confocal fluorescent images of microtubules (green) and the edges of the ER accumulation around spindles (white) in an ascidian embryo from 1-cell stage metaphase to 4-cell stage interphase. Ant. is anterior, Veg. is vegetal, and NEB is nuclear envelope breakdown. Time is in minutes:seconds. Scale bar, 30 µm. **(B)** Plot of the rotation and tilt angles between the CCs or the spindles of the left and right cells at the 2-cell stage as a function of time in control (*N* = 9 embryos) and in calcium and magnesium-free seawater (CMF)-treated embryos (*N* = 10 embryos). The dashed line at 17 min indicates NEB. Error bars, standard error of the mean. **(C)** Plot of the initial (Centrosome separation) and final (Metaphase 2-cell) rotation (left) and tilt (right) angles between the left and right CCs or spindles at 2-cell stage in control embryos (*N* = 9 embryos; rotation: paired Wilcoxon test, *P=0.0131; tilt: paired Wilcoxon test, *P=0.0250) and CMF-treated embryos (*N* = 10 embryos; rotation: paired t test, ns - not significant; tilt: paired t test, ns – not significant). Control vs. CMF rotation angle: one-way ANOVA (***P=0.0010) with Šídák’s multiple comparisons test (ns – not significant, *P=0.0431). Control vs. CMF tilt angle: Kruskal-Wallis (ns – not significant) with Dunn’s multiple comparisons test ns – not significant).

**Figure S5:**
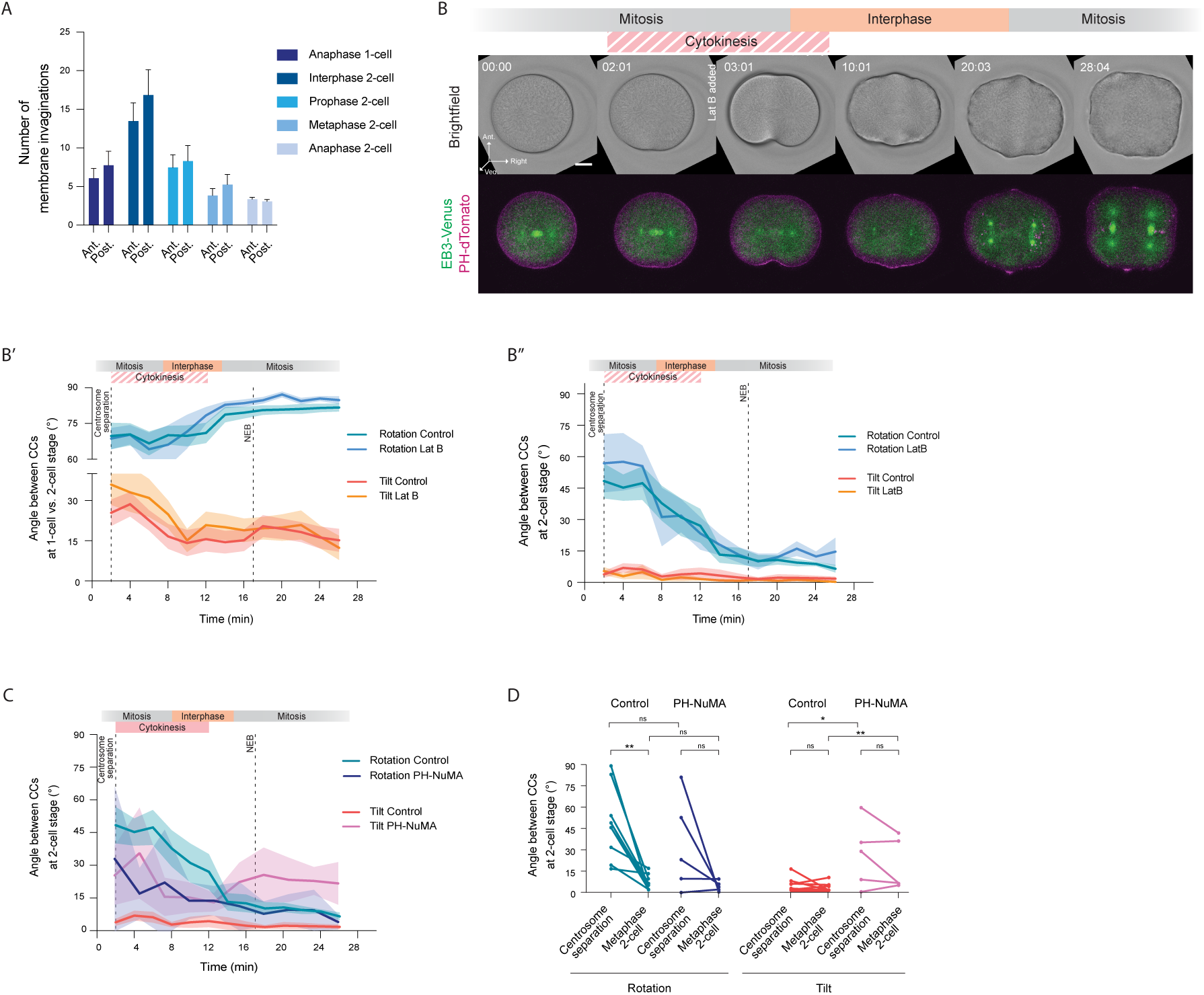
Cortical pulling is increased in the posterior of the ascidian 2-cell stage embryo (relative to Fig. 5). **(A)** Bar plot of the average number of membrane invaginations at the anterior (Ant.) and posterior (Post.) sides of the embryo as a function of the cell cycle stage. Data is taken from Fig. 5B’. **(B)** Brightfield and confocal fluorescent images of a central z-slice of the plasma membrane (magenta), and maximum intensity projections of centrosomes (green) in an ascidian embryo from 1-cell stage metaphase to 2-cell stage metaphase treated with 1 µg/ml Latrunculin B at 1-cell stage anaphase. Ant. is anterior, Veg. is vegetal. Time is in minutes:seconds. Scale bar, 30 µm. **(B’)** Plot of the rotation and tilt angles between the CCs or the spindles at the 2-cell stage and the spindle at the 1-cell stage as a function of time in control embryos (*N* = 9 embryos, *n* = 18 cells) and Latrunculin B-treated embryos (*N* = 5 embryos, *n* = 10 cells). The dashed line at 17 min indicates NEB. Error bars, standard error of the mean. **(B’’)** Plot of the rotation and tilt angles between the CCs or the spindles of the left and right cells at the 2-cell stage as a function of time in control (*N* = 9 embryos) and Latrunculin B-treated (*N* = 5 embryos). The dashed line at 17 min indicates NEB. Error bars, standard error of the mean. **(C)** Plot of the rotation and tilt angles between the CCs or the spindles of the left and right cells at the 2-cell stage as a function of time in control (*N* = 9 embryos) and embryos expressing PH-NuMA (*N* = 5 embryos, *n* = 10 cells). The dashed line at 17 min indicates NEB. Error bars, standard error of the mean. **(D)** Plot of the initial (Centrosome separation) and final (Metaphase 2-cell) rotation (left) and tilt (right) angles between the left and right CCs or spindles at 2-cell stage in control embryos (*N* = 9 embryos; rotation: paired Wilcoxon test, *P=0.0131; tilt: paired Wilcoxon test, *P=0.0250) and embryos overexpressing PH-NuMA (*N* = 5 embryos; rotation: paired t test, ns - not significant; tilt: Wilcoxon test, ns – not significant). Control vs. PH-NuMA rotation angle: one-way ANOVA (***P=0.0005) with Šídák’s multiple comparisons test (ns – not significant). Control vs. PH-NuMA tilt angle: Kruskal-Wallis test (*P=0.0210) with Dunn’s multiple comparisons test (*P=0.0146, ns – not significant).

## MOVIE LEGENDS

**Movie 1.** Centrosomal complex orientation during the first two cleavages in ascidian embryos, related to Fig. 1 and S1. Time-lapse brightfield (left panel) and fluorescent confocal video of a representative ascidian embryo injected with PH-dTomato mRNA (magenta, right panel) and EB3-3xVenus mRNA (green, right panel) to mark membrane and centrosomes, respectively. Time is in minutes:seconds. Time interval, 2 min.

**Movie 2.** Centrosome and nucleus migration during interphase of the 2-cell stage in ascidian embryos, related to Fig. 2. Time-lapse fluorescent confocal video of a representative ascidian embryo injected with EB3-mScarlet mRNA (green), H2B-Venus mRNA (magenta) to mark centrosomes and the DNA, respectively, and incubated in MitoTracker^TM^ Deep Red (cyan) to label the mitochondria. Time is in minutes:seconds. Time interval, 16 s.

**Movie 3.** Laser ablation of intercentrosomal microtubules during CC migration at the 2-cell stage of ascidian embryos, related to Fig. 2 and S2. Time-lapse fluorescent confocal video of a representative ascidian embryo injected with EB3-mScarlet mRNA (magenta) and Ensconsin-3xGFP mRNA (green) to mark centrosomes and microtubules, respectively. The ablation site is indicated by the arrowhead. Time is in minutes:seconds. Time interval, 12.8 s.

**Movie 4.** Laser ablation of microtubules between the CC and the nucleus during CC migration at the 2-cell stage of ascidian embryos, related to Fig. 2 and S2. Time-lapse fluorescent confocal video of a representative ascidian embryo injected with H2B-Venus mRNA (magenta) and Ensconsin-dTomato mRNA (green) to mark DNA and microtubules, respectively. The ablation site is indicated by the arrowheaad. Time is in minutes:seconds. Time interval, 12.8 s.

**Movie 5.** Laser ablation of astral microtubules during CC migration at the 2-cell stage of ascidian embryos, related to Fig. 3 and S3. Time-lapse fluorescent confocal video of a representative ascidian embryo injected with EB3-mScarlet mRNA (magenta) and Ensconsin-3xGFP mRNA (green) to mark centrosomes and microtubules, respectively. The ablation site is indicated by the arrowhead. Time is in minutes:seconds. Time interval, 12.8 s.

**Movie 6.** Endoplasmic reticulum rearrangement during the first two cleavages in ascidian embryos, related to Fig. 4 and S4. Time-lapse fluorescent confocal video of a representative ascidian embryo injected with PH-dTomato mRNA (magenta) and Reticulon-Venus mRNA (grey) to mark the membrane and endoplasmic reticulum, respectively. Time is in minutes:seconds. Time interval, 110 s.

**Movie 7.** Centrosomal complex orientation during the first two cleavages in ascidian embryos cultured in calcium and magnesium-free seawater, related to Fig. 4 and S4. Time-lapse brightfield (left panel) and fluorescent confocal video of a representative ascidian embryo injected with PH-dTomato mRNA (magenta, right panel) and EB3-3xVenus mRNA (green, right panel) to mark membrane and centrosomes, respectively. Time is in minutes:seconds. Time interval, 2 min.

**Movie 8.** Cortical pulling assay in control ascidian embryos, related to Fig. 5 and S5. Time-lapse fluorescent confocal video of a representative ascidian embryo treated with 1μg/ml Latrunculin B and injected with H2B-Venus mRNA (magenta) to mark the DNA, and incubated in CellMask^TM^ Orange to label the plasma membrane (grey). Time is in minutes:seconds. Time interval, 20 s.

**Movie 9.** Cortical pulling assay in ascidian embryos expressing PH-NuMA, related to Fig. 5 and S5. Time-lapse fluorescent confocal video of a representative ascidian embryo treated with 1μg/ml Latrunculin B and incubated in CellMask^TM^ Orange to label the plasma membrane (grey, left panel) and injected with PH-NuMA-Venus mRNA (magenta, right panel) to mark exogenous cortical NuMA. Time is in minutes:seconds. Time interval, 16 s.

**Movie 10.** Centrosomal complex orientation during the first two cleavages in ascidian embryos expressing PH-NuMA to increase cortical pulling, related to Fig. 5 and S5. Time-lapse brightfield (left panel) and fluorescent confocal video of a representative ascidian embryo injected with PH-NuMA-Venus mRNA (magenta, right panel) and EB3-mScarlet (green, right panel) to mark exogenous cortical NuMA and centrosomes, respectively. Time is in minutes:seconds. Time interval, 2 min.

